# MDF regulates a network of auxin-dependent and -independent pathways of adventitious root regeneration in *Arabidopsis*

**DOI:** 10.1101/2024.05.26.595954

**Authors:** Fahad Aldowigh, Rodrigo Matus, Julien Agneessens, Haozhan Gao, Wenbin Wei, Jennifer Topping, Keith Lindsey

**Affiliations:** Department of Biosciences, Durham University, Stockton Road, Durham DH1 3LE, UK

**Author notes:** Biology Department, College of Applied Sciences, Umm al-Qura University, Makkah 24381, Saudi Arabia (Fahad Aldowigh). Azotic Technologies Limited, Chessingham Park, York YO19 5SN, UK.

## Abstract

Plants exhibit strong plasticity in growth and development, seen clearly in lateral and adventitious root development from differentiated tissues in response to environmental stresses. Previous studies have demonstrated the role of both auxin-dependent and auxin-independent signalling pathways in regulating the *de novo* formation of adventitious roots (ARs) from differentiated tissues, such as leaf petiole in *Arabidopsis.* One important question is how the auxin-dependent and -independent pathways are coordinated. To investigate this question, we used a combined approach of inducible gene expression, mutant, and signalling reporter gene analysis during AR regeneration in the *Arabidopsis* petiole to understand regulatory relationships. Auxin signalling components AXR1 and AXR3, and the PIN trafficking protein VAMP714, are each required for AR initiation, as is the ethylene signalling repressor POLARIS, but not EIN2. We identify the RNA splicing regulator MDF and the transcription factor RAP2.7 as new positive regulators of both the auxin-independent and auxin-dependent pathways, and show that MDF regulates *RAP2.7*, *WOX5* and *NAC1*; while RAP2.7 regulates *WOX5* but not *NAC1* or *YUC1*. NAC1 is required for *de novo* root formation in a pathway independent of *YUC1*, *WOX5* or *RAP2.7*. We propose a model in which MDF represents a point of molecular crosstalk between auxin-dependent and -independent regeneration processes.

## Introduction

An important question in developmental biology is that of how new cell identities are acquired during organogenesis. In plants, regeneration of new organs following re-specification of cell identity is more common than in animals, but the mechanisms are incompletely understood (Xu and Huang, 2014; Jing et al., 2020). Adventitious roots (ARs) represent one form of regeneration from previously differentiated cells. They are typically initiated in response to changes in the environment such as flooding, drought, or wounding (Steffens and Rasmussen, 2016), and help plants to acclimate to stressful environments. For example, ARs help plants absorb water and oxygen in flooding, and they can grow from wound sites as part of a regeneration process (Gonin et al., 2019). ARs that develop from a wound site represent *de novo* organogenesis. This kind of root developmental programme depends on endogenous hormones and is considered to arise from procambium or cambium cells, which contain stem cells located in the vascular tissues of aerial organs (Chen et al., 2014; Liu et al., 2014).

ARs can be grown from non-root tissue in isolation in response to hormonal signals, notably auxin. There are two promoting principles involved in damage-related formation of ARs (Chen et al., 2014). The first is the wound response that is induced during the isolation of the explant tissue from resource supply in the plant. The second is the signalling networks that affect organ regeneration post-wounding. For example, detached leaf explants can activate various early signals that include both short- and long-range signals that lead to AR formation (Chen et al., 2014). AR formation from leaf explants has been considered to comprise three developmental phases: the early wound response, the creation of an auxin maximum in competent cells and the establishment of founder cells, leading to *de novo* root meristem formation and activity (Jing et al., 2020). Wound signals induce *YUCCA1* (*YUC1*) and *YUC4* expression near the wound region to help maintain auxin levels (Xu, 2018). YUC1 and YUC4 mediate auxin biosynthesis and they are important for cell fate transition during *de novo* organogenesis (Chen et al., 2016a, b). The YUC family also plays an important role in converting indole-3-pyruvic acid (IPA) to indole-3-acetic acid (IAA) (Sun et al., 2016). Cell fate transition is blocked if auxin signalling is inhibited (Liu et al., 2014; Xu, 2018).

Wound signals have many functions, including the activation of expression of the *NAC1* gene (Xu, 2018). NAC1 (NAC DOMAIN-CONTAINING PROTEIN 1) is a transcription factor of the NAC family required for *de novo* root formation, and transgenic repression of *NAC1* expression causes reduced development of adventitious roots (Chen et al., 2016c). The NAC pathway works independently from the auxin pathway and wounding plays an important role in inducing it in leaf explants of *Arabidopsis*. In addition, NAC1 may have a role in cell wall metabolism to promote AR development (Chen et al., 2016c). It is also suggested that *NAC* genes promote expression of KDEL-tailed Cys endopeptidase (CEP) genes (*CEP1* and *CEP2*). CEP proteins have been found to contribute to programmed cell death, by their secretion to the cell wall to induce inactivation of EXTENSIN (EXT) proteins. EXT proteins are important for cell wall expansion and wound healing, and wound healing may antagonise AR emergence. Therefore, repression of *EXT* genes by NAC proteins may promote AR emergence from leaf explants (Chen et al., 2016c). *NAC1* appears to be working independently of the auxin pathway and may have a role in cell wall metabolism to promote AR development. However, the regulation of *NAC1* in the context of *de novo* root regeneration is not well understood.

The third phase of AR formation, namely the fate determination of founder cells and their subsequent development into a functional root meristem, can be described as involving four steps (Jing et al., 2020). The first step of cell fate determination (’Priming’) involves auxin activation of the expression of *WUSCHEL RELATED HOMEOBOX11* and *12* (*WOX11/12*) genes, involved in the transformation of competent cells to root founder cells in the leaf explants. The second step (’Initiation’) involves cell division to create root primordium cell layers around 2-4 days after wounding (Xu, 2018). The homeobox gene *WOX5* is required to start this second step whereby root founder cells divide to become root primordium cells. In addition, transcriptional activation of *WOX5* requires auxin, and repression of auxin causes reduced expression of *WOX5* which in turn reduces *de novo* root formation (Hu and Xu, 2016). The third step (’Patterning’) creates cellular pattern in the root apical meristem and is characterized by the loss of *LATERAL ORGAN BOUNDARIES DOMAIN16* (*LBD16*) expression and restriction of *WOX5/7* expression to the stem cell niche. Other regulatory genes such as *SHORT ROOT* (*SHR*), *PLETHORA1/2* and *SCARECROW* (*SCR*) are also involved in meristem formation (Bustillo-Avenado et al., 2018; Kim et al., 2018). Finally, the fourth step (’Emergence’) is characterized by root tip growth through the epidermis of the leaf explant.

The aim of the work described in this paper was to understand better the mechanistic basis and inter-relationship between the auxin-dependent and -independent control of *de novo* root regeneration. We show a requirement for the auxin-regulated R-SNARE VESICLE-ASSOCIATED MEMBRANE PROTEIN 714 (VAMP714) in AR formation and growth, and identify the RNA splicing regulator MERISTEM-DEFECTIVE (MDF; Casson et al., 2009; Thompson et al., 2023) and the transcription factor RAP2.7 (RELATED TO AP2.7; Tair 2024) as new positive regulators of both the auxin-independent pathway and auxin-dependent pathways. We then demonstrate that MDF regulates *RAP2.7*, *WOX5* and *NAC1*; while RAP2.7 regulates *WOX5* but not *NAC1* or *YUC1*. NAC1 is required for *de novo* root formation in a pathway independent of *YUC1*, *WOX5* or *RAP2.7*. We propose a model for the network of gene and hormone signalling interactions involved in adventitious root formation and suggest that MDF represents a point of molecular crosstalk between auxin-dependent and -independent regeneration processes.

## Results

### Contrasting roles of AXR1 and AXR3 and ethylene signalling in *de novo* root initiation and growth

A simple method to induce the *de novo* development of roots *in vitro* has been described by Chen et al. (2014). This involves wounding the petiole of a 12 d-old leaf of *Arabidopsis*, and then incubating the leaf on B5 nutrient medium lacking both hormones and sucrose. New meristem initiation from the wild type petiole wound site is evident by ca. 5 d after wounding (DAW), associated with an auxin maximum as indicated by expression of the auxin reporter *DR5::GUS* in the new root primordium; and *de novo* roots emerging after 8-10 DAW (Suppl. Fig. S1). To gain new insight into the roles of specific signalling pathway regulators during *de novo* root meristem formation and growth, 12 d-old leaves were wounded by excision from the seedling of signalling mutant genotypes, and the root regeneration and growth responses compared to wild type responses were determined. The focus of this study was the interrelationship between auxin and ethylene pathways and associated transcriptional and signalling regulators with likely roles in meristem function in seedlings of Arabidopsis.

Fig. 1A shows the mean total numbers of root branches (*de novo* adventitious roots up to 12 DAW, plus associated lateral roots derived from them for subsequent days) formed from wounded petioles of wild type (Col-0) and mutants defective in auxin signalling (auxin-resistant *axr1* and auxin-hypersensitive *axr3*; Lincoln et al., 1990; Leyser et al., 1996) and ethylene signalling (ethylene-insensitive *ein2*, ethylene hypersignalling *pls*; Casson et al., 2002, Chilley et al., 2006) after 12, 19 and 26 DAW. For wild type, the mean number was 1.4 (n > 30) at 12 DAW, rising to 36.8 at 26 DAW. *axr1* was unable to regenerate any roots, while *axr3* produced a mean of 2.1 at 12 DAW (not significantly different to wild type), rising to 13.7 (i.e. ca. 37% that of wild type, significantly less than wild type, P <0.01) at 26 DAW. The *ein2* leaf produced a mean of 1.3 branches at 12 DAW, not significantly different to wild type either then or at 19 DAW, but produced a significantly lower number of branches at 26 DAW, with 23.5 (ca. 64% of wild type; P <0.05). *pls* produced a mean of 0.9 root branches at 12 DAW, rising to 3.1 at 26 DAW (ca. 12% of wild type; P <0.001).

**Figure 1.**
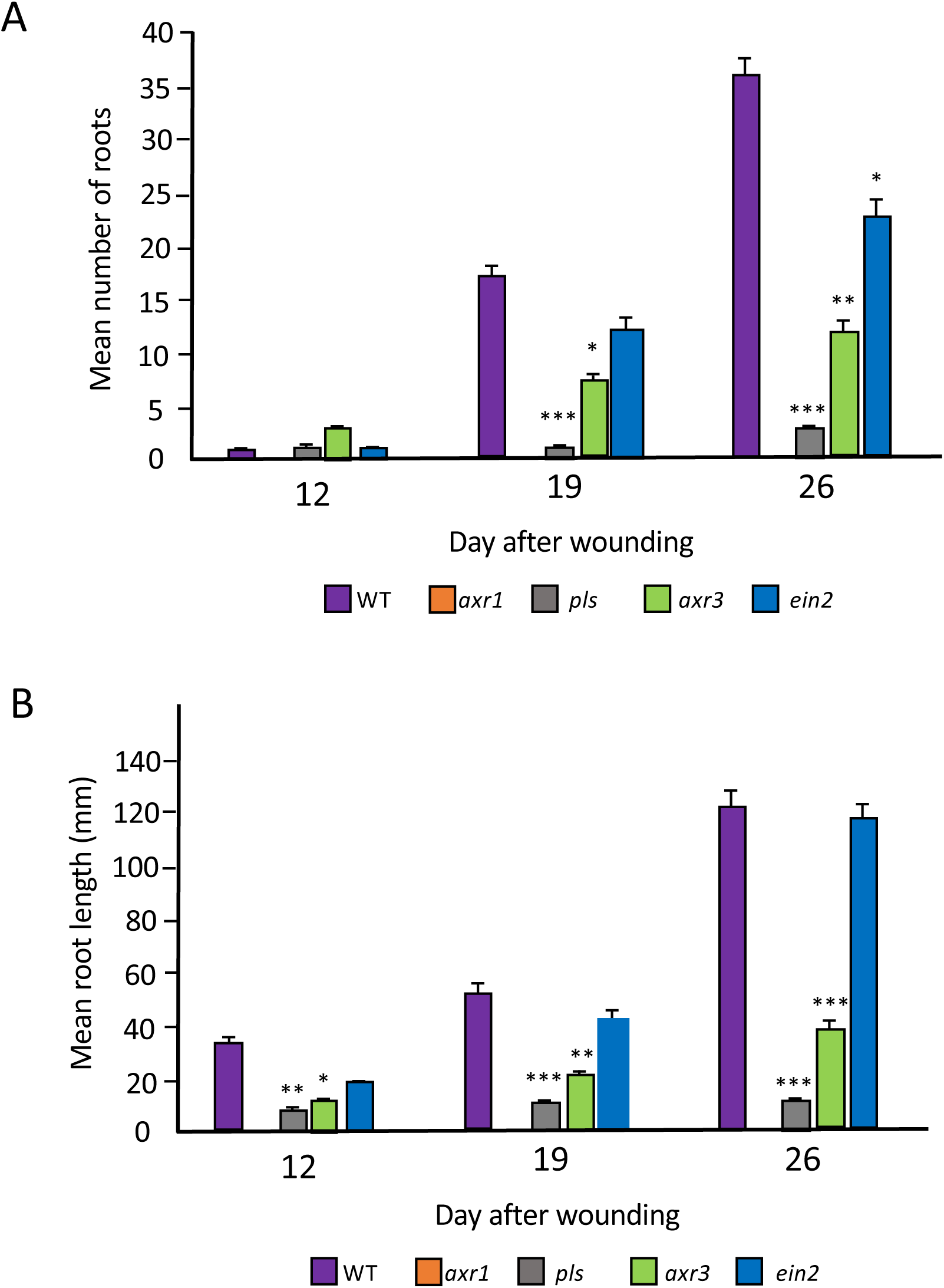
(A) Mean number of regenerated roots and (B) mean root lengths for wild type and mutants at 12, 19 and 26 d after wounding on B5 medium. Values are means of at least 30 samples + SEM. Statistical significance was determined using Student’s t-test for independent samples compared to wild type values, with P-values <0.05 (*), P <0.01 (**), P-value < 0.001 (***).

Fig. 1B shows the mean lengths of the primary adventitious roots initiated from wounded petioles of wild type and the auxin and ethylene signalling mutants. By 12 DAW the mean primary root length for Col-0 was 34.25 mm. For *axr1* no roots were detected at 12 DAW, for *axr3* the mean length was 11.49 mm, for *ein2* 20.19 mm, and for *pls* 6.51 mm. By 19 DAW, the wild type mean root length increased to 52.2 mm, *axr3* roots increased to a mean of 23.17 mm, *pls* increased by a much smaller amount to a mean of 7.75 mm, and *ein2* roots were not significantly different to wild type at a mean of 43.5 mm. This general pattern was retained at 26 DAW (Col-0 122.41 mm, *axr3* 39.19 mm, *ein2* 117.61 mm, *pls* 8.30 mm, with no roots on *axr1*).

These results suggest AXR1-mediated auxin signalling, but not AXR3-mediated auxin or PLS- or EIN2-mediated ethylene signalling, is essential for root initiation, given the number of roots produced by 12 DAW is similar in the *axr3*, *pls* and *ein2* mutants. However, AXR3 and possibly also AXR1 (both required for correct auxin responses) and PLS (which represses ethylene responses) are required for wild type levels of root elongation and lateral root initiation from primary adventitious roots.

### Hormone signalling gene interactions in regeneration control

Previous work has shown a requirement for auxin biosynthesis for *de novo* root regeneration, via the YUCCA pathway (Chen et al., 2016a, b). Matosevich et al. (2020) suggested that, in contrast to earlier views (Sena et al., 2009), PIN-dependent polar auxin transport is not essential for root tip regeneration from wounded roots, based on analysis of *pin* mutants and naphthylphthalamic acid (NPA) treatments (NPA inhibits PIN protein activity; Abas et al., 2021). This is indicative of an auxin-independent pathway for *de novo* root regeneration, though there was suggested a possible role for short-distance (symplastic) auxin transport at the wound surface (Mellor et al., 2020). Nevertheless, *PIN1*, *PIN3*, *PIN7* and auxin-inducible *DR5::GUS* expression is induced in the leaf petiole, in particular during regeneration (Suppl. Figs. S1-4), suggesting that the polar auxin transport pathway is activated. VAMP714 is an R-SNARE vesicle-associated protein required for the trafficking of PIN proteins and polar auxin transport (Gu et al., 2021). The *VAMP714::GUS* promoter-reporter is auxin-regulated and is expressed in a similar pattern to *DR5::GUS* (Suppl. Fig. S5). The *vamp714* loss-of-function mutant, defective in polar auxin transport, shows similar adventitious root initiation compared to wild type (Fig. 2A, 12 and 17 DAW) but reduced lateral root initiation and growth by 21 DAW (Fig. 2B), suggesting VAMP714-mediated PIN localization is not required for AR initiation, but is required for later stages of root development from ARs. Loss of VAMP714 activity is associated reduced expression of auxin-regulated gene *IAA2* in both explanted leaf blade and petiole of the mutant (Fig. 2, C). In any case a new auxin maximum is established close to the wound site. The synthetic *EBS::GUS* ethylene reporter was expressed in the leaf blade up to 8 DAW, and in the petiole but not root primordium at 5-6 DAW (Suppl Fig. S6). However, after 8 DAW GUS expression occurred in the emerging *de novo* root, suggesting ethylene responses are not obviously important in early stages of primordium development; though the *pls* mutant data (Fig. 1A) indicate there may be required a regulated ethylene signalling response for adventitious root elongation.

**Figure 2.**
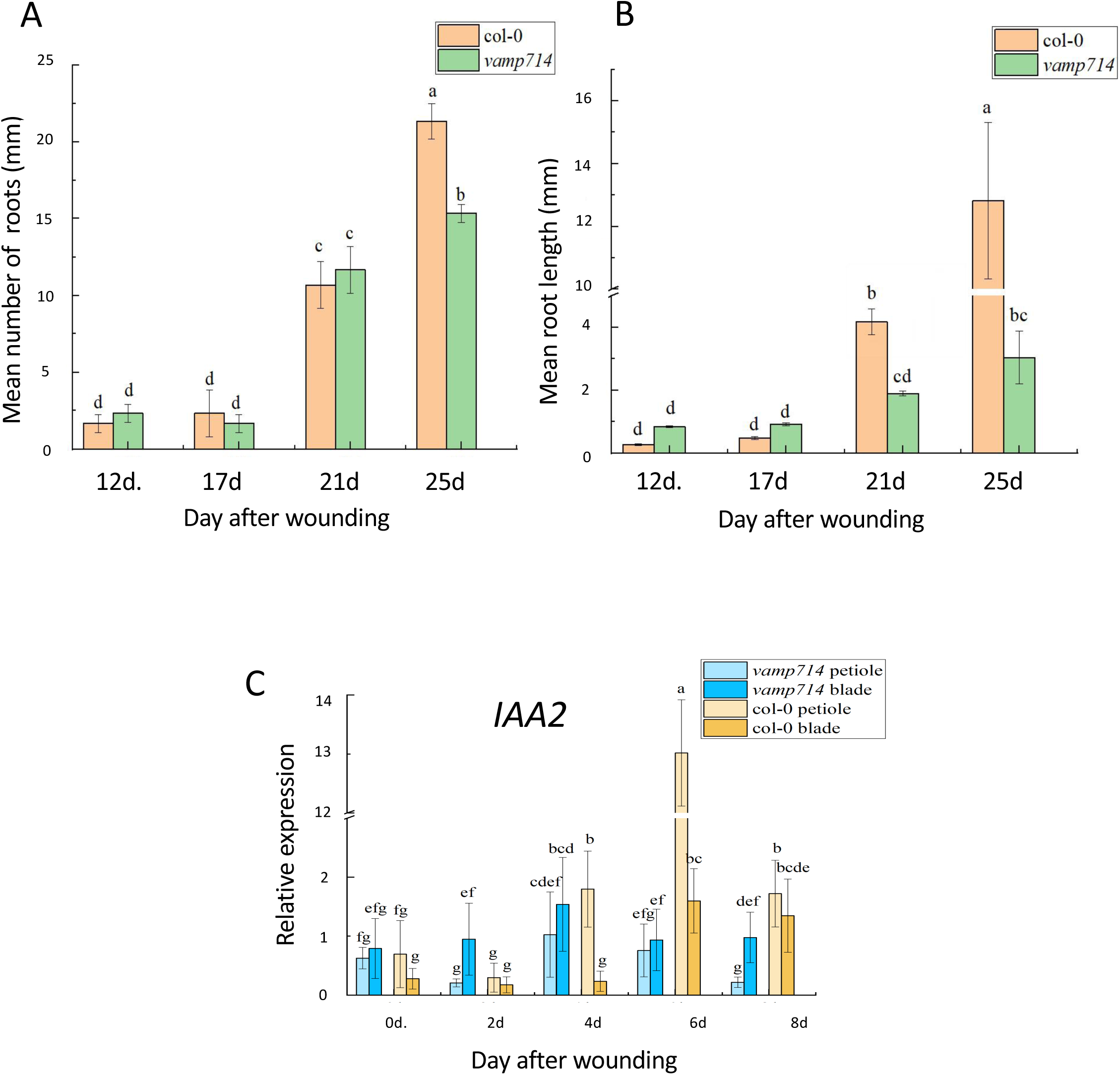
(A) Mean number of regenerated roots and (B) mean root lengths for wild type (Col-0) and loss-of-function *vamp714* mutant at 12 d, 17d, 21d, 25d after wounding on B5 medium. (C) RT-qPCR analysis of *IAA2* gene expression in wild type and mutant (*vamp714*) leaf blade and petiole at 0 d, 2 d, 4 d, 6 d and 8 d after wounding, using *ACTIN2* as reference gene. Values represents means and error bars are SEM (n = three biological repeats with three technical repeats). One-way ANOVA and correlation analysis was performed using SPSS 26.0 software, and Duncan ’s new multiple range method was used for significance test (P < 0.05 ).

To determine the relationship between hormone signalling and regulatory genes known to be involved in root regeneration, we first used RT-qPCR to measure the expression of the genes *NAC1, WOX5*, *YUC1* and *YUC4* in the auxin and ethylene signalling mutants *axr1*, *axr3*, *pls* and *ein2* compared to expression in wild type during a root regeneration time course of 0, 2, 3, 7 and 14 DAW. Fig. 3A shows a significant increase in *NAC1* expression between 2 and 3 DAW in all genotypes, with no significant difference between wild type and mutants during this period. Interestingly, by 14 DAW *NAC1* expression in wild type, *ein2* and *axr3* has continued to increase, but has dramatically reduced in both *pls* and *axr1* mutants, associated with reduced (in *pls*) or no (in *axr1*) root initiation and growth in these mutants by this stage. It is notable however that during the period in which *de novo* root regeneration is initiated (5-8 DAW), *NAC1* expression is unaffected in ethylene or auxin signalling mutants compared to expression levels in wild type.

**Figure 3.**
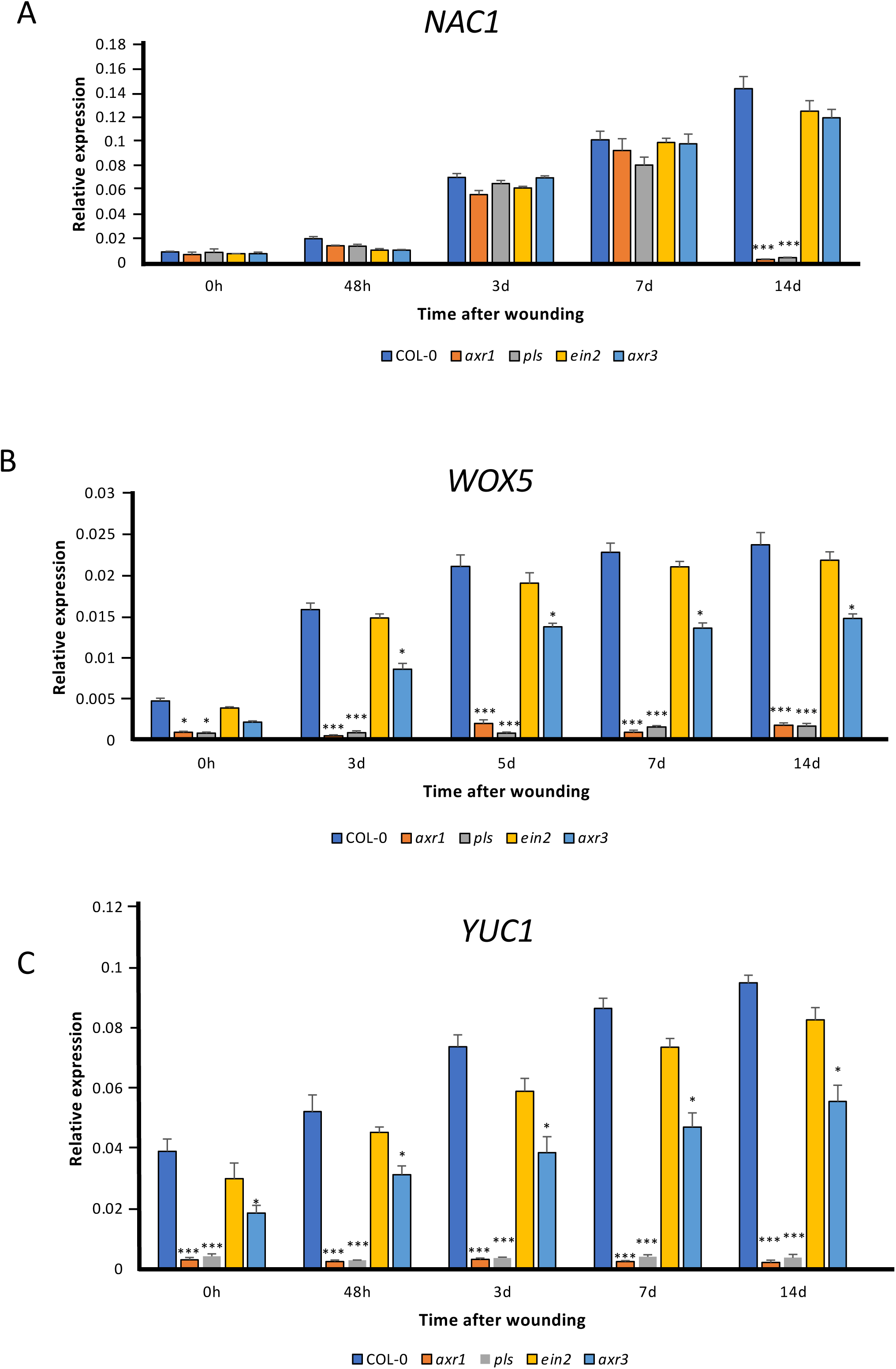
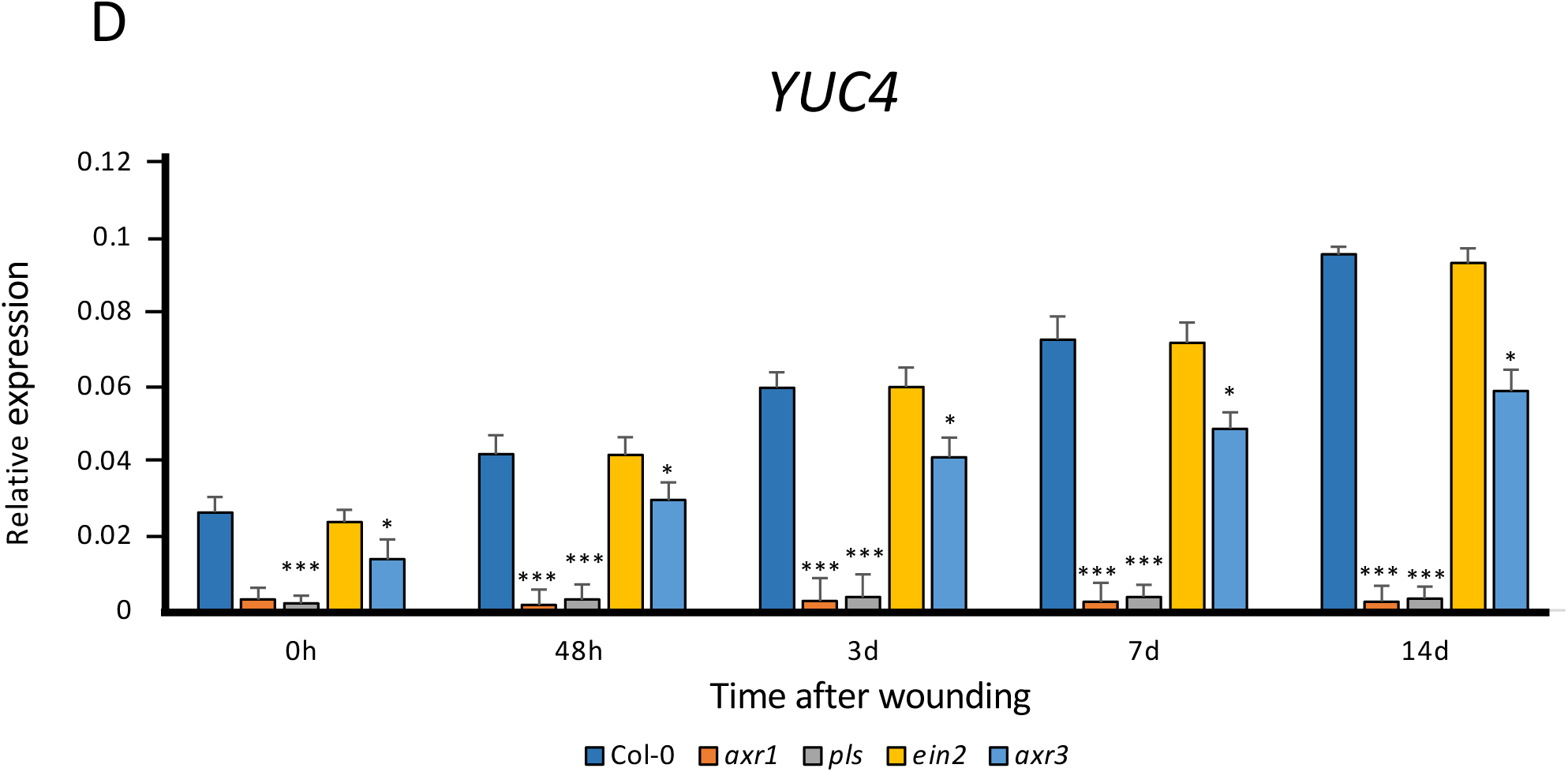
RT-qPCR analysis of (A) *NAC1* (B) *WOX5* (C) *YUC1* and (D) *YUC4* gene expression in wild type and mutant leaf (*axr1*, *pls*, *ein2* and *axr3*) at 0 h, 48 h, 3 d, 7 d and 14 d after wounding, using *UBC* as reference gene. Values represents means and error bars are SEM (n = three biological repeats with three technical repeats). Statistical significance was determined using Student’s t-test for independent samples compared to wild type values, with P-values <0.05 (*), P <0.01 (**), P-value < 0.001 (***).

Fig. 3, B-D show there are very similar patterns of expression of each of *WOX5*, *YUC1* and *YUC4* during the culture and regeneration period, in contrast with the pattern for *NAC1*. *WOX5*, *YUC1* and *YUC4* showed increased expression in wild type, *ein2* and *axr3* samples during the culture period, but very low levels of expression were seen in *axr1* and *pls* from the beginning of the experiment, with no subsequent increase. For *axr3*, the expression of each gene was significantly lower than in wild type and *ein2* from 3 DAW onwards, though significantly higher than in either *axr1* or *pls*, which showed the lowest levels of expression of *WOX5*, *YUC1* and *YUC4*. These results indicate that *WOX5*, *YUC1* and *YUC4* are dependent on both AXR1 and PLS for expression during *de novo* root formation. *NAC1* expression was not dependent on the auxin signalling *AXR1* and *AXR3* genes, nor the ethylene signalling *PLS* and *EIN2* genes during the regeneration process (4-8 DAW).

### MDF regulates both auxin-dependent and -independent root regeneration

Next we investigated the possible regulatory role in *de novo* regeneration of three transcriptional regulatory genes with roles in meristem function (*MDF*, *RAP2.7* and *NAC1)*. The *MDF* gene of Arabidopsis is required for root meristem organization and maintenance (Casson et al., 2009). It encodes a putative RS domain protein and has recently been demonstrated to be a splicing factor that regulates meristem function through both auxin-dependent and auxin-independent mechanisms, to maintain stemness (Thompson et al., 2023). Our analysis of the RNA-seq data produced for the *de novo* regeneration of roots from leaf petioles by Liu et al. (2022) shows that *MDF* is expressed throughout a 5 d culture and regeneration period, with a peak in expression at around 3 DAW (Suppl. Fig. 7, A). Its expression is independent of auxin transport as indicated by no clear change in level or pattern across the time course in the presence of NPA compared to in the absence of NPA (Suppl. Fig. S7, B). This auxin independence is consistent with *MDF* regulation in intact seedlings (Casson et al., 2009). However there is evidence that MDF regulates downstream auxin-related pathways. The *mdf* mutant shows decreased levels of *PIN* family mRNAs and this is associated with a reduced auxin maximum in the basal region of the *mdf* embryo and seedling root meristem. Furthermore, seedling roots of *mdf* show reduced expression of *SHORTROOT*, *SCARECROW*, *WOX5* and *PLETHORA* genes (Casson et al., 2009, Thompson et al., 2023). RNA-seq analysis shows slightly reduced *VAMP714* expression in *mdf* (log2-fold 0.74, padj = 6.26E-10; Thompson et al., 2023), which may contribute to the observed defective PIN localization in the *mdf* mutant.

The *RAP2.7* gene is a member of the AP2/ERF family of transcription factors which are involved in regulating the process of flowering and innate immunity and may be involved in the control of meristematic activity by an as yet unknown mechanism (Tair, 2024). We recently found *RAP2.7* to be a splicing target of MDF (it is mis-spliced in the *mdf* mutant; Thompson et al., 2023) and has also been identified as being expressed in the *Arabidopsis* root tip, similar to *MDF* (Birnbaum et al., 2003). The RNA-seq data of Liu et al. (2022) show that, like *MDF*, *RAP2.7* expression is detectable throughout a 5 day leaf culture and AR regeneration period, with a peak at ca. 12 hours after wounding (HAW); and, similar to *MDF*, its pattern and level of expression is largely unaffected by NPA treatment, indicative of it not being auxin-regulated (Suppl. Fig. S7, C and D).

Given that the *NAC1* gene is known to be involved in *de novo* root formation (Chen et al., 2016c) and in lateral root development mediated by auxin signalling (Xie et al., 2000), we were interested in understanding its functional relationship, if any, with *MDF* and *RAP2.7*.

For analysis of *MDF* function, we utilized the previously described *mdf-1* loss-of-function mutant, and also generated transgenic plants containing an estradiol-inducible *XVE-35S::MDF* gene construct for overexpression analysis. Three independent transformants were characterized, all showing ca. 6-7-fold induction of *MDF* expression, and one representative line was chosen for further analysis (Suppl. Fig. S8). The results in Fig. 4, A and B show the numbers and lengths of *de novo* roots (including associated lateral roots) formed from leaf of wild type, *MDF* transgenic overexpresser (MDF-OV) following estradiol induction, and the *mdf-1* mutant at 12, 19 and 26 DAW. By 12 DAW the mean number of roots was 1.66 for Col-0 and 2.56 for MDF-OV, which was not statistically significantly different. By 19 DAW the number of roots increased (mean of 24.2 roots for Col-0, 32.44 branches for MDF-OV) and there was a statistically significant difference between Col-0 (mean 40.1 roots initiated) and MDF-OV (mean 74.8 roots initiated; ca. 187% of wild type; P <0.01, n = 30) by 26 DAW. The *mdf* mutant failed to produce any detectable roots over the entire time course. This was associated with reduced expression of the auxin-responsive *IAA1* gene over the 8 d regeneration time course (Suppl. Fig. S9). At 12 DAW, the roots produced from Col-0 and MDF-OV leaves showed no statistically significant difference in mean length; but at both 19 and 26 DAW the *MDF* overexpresser produced significantly longer roots than wild type (P < 0.01, n = 30; Fig. 4B). This shows a positive regulatory role for MDF in both initiation and growth of *de novo* adventitious roots in this system.

**Figure 4.**
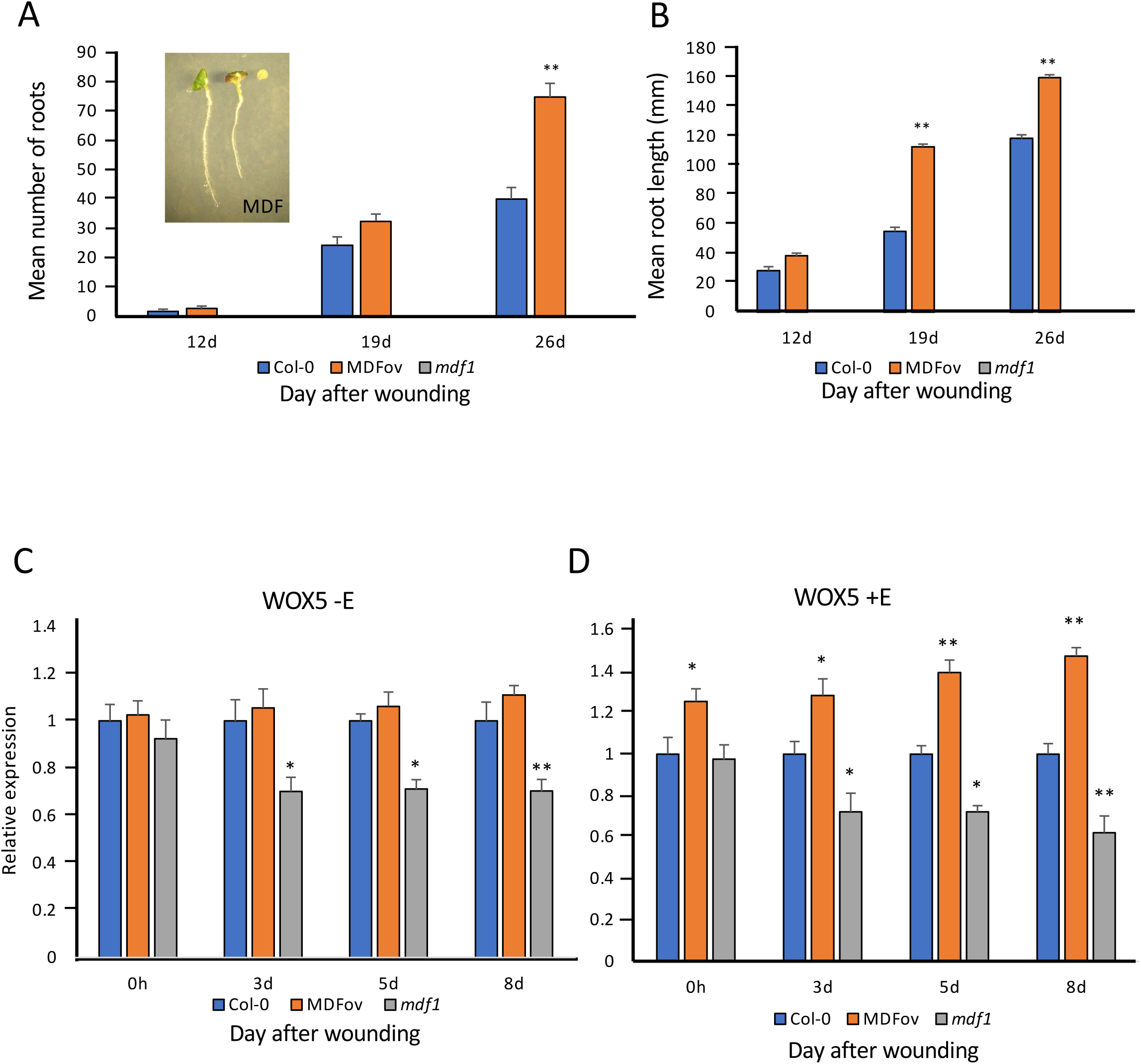
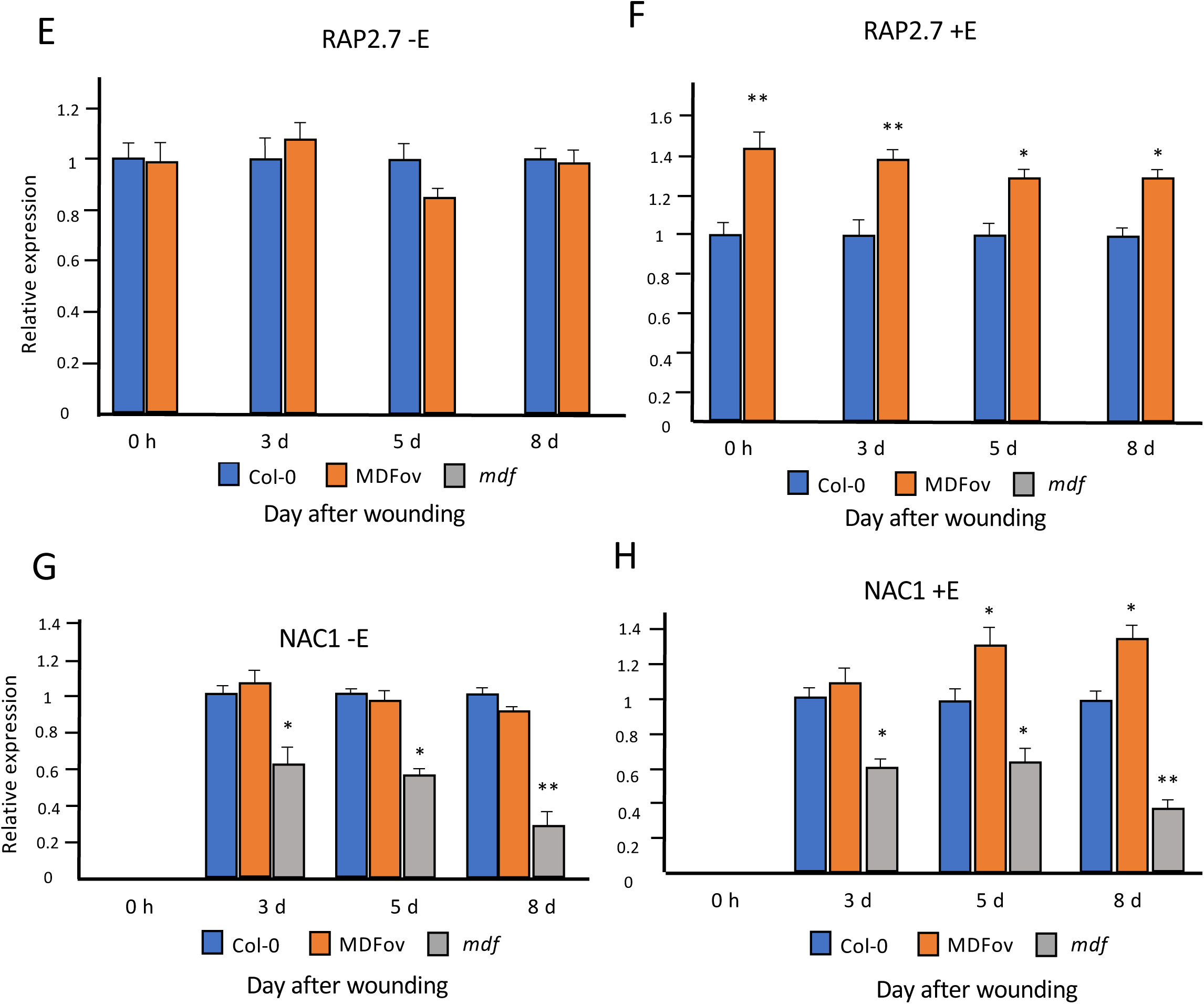
(A) Mean numbers of regenerated roots in wild type Col-0, estradiol-induced *MDF* overexpression (MDFov) and *mdf-1* after estradiol treatment and wounding on B5 medium for 12, 19 and 26 d. Inset: (left) MDFov leaf with *de novo* root after 12 d; (centre) Col-0 leaf with *de novo* root after 12 d; (right) *mdf-1* leaf after 12 d; (B) Mean root length in wild type Col-0, MDFov and *mdf-1* after estradiol treatment and culture on B5 medium for 12, 19 and 26 d. Values are averages of at least 30 leaves ± SEM. Statistical significance (A,B) was determined using Student’s t-test for independent samples compared to wild type values, with P-values P<0.01 (**). Scale bar = 1 cm. (C-H). Relative expression of *WOX5*, *RAP2.7* and *NAC1* genes in wild type (Col-0), MDFov and *mdf-1* samples at 0 h, 3 d, 5d and 8 d after germination on B5 medium either with estradiol treatment (+E) or without (-E) estradiol treatment to induce *MDF* overexpression (MDFov). Values are means ± SEM for three biological repeats. Statistical significance was determined using Student’s t-test for independent samples compared to wild type values, with P-values P< 0.05 (*) and P<0.01 (**).

To determine whether MDF has a role in the regulation of the *WOX5* gene, which is shown in Fig. 3B to be associated with adventitious root regeneration, RNA was extracted from cultured leaf of wild type, *mdf-1* and the MDF-OV line, either untreated (-E) or treated with estradiol (+E) to induce *MDF* expression over an 8 d time course of culture, and transcriptional analysis was carried out. Fig. 4, C and D show the mean expression data from three biological replicate experiments, each with three technical replicates. In the absence of estradiol induction, there was no significant difference in *WOX5* expression between wild type (Col-0) and MDF-OV samples, but there was a statistically significantly lower level of expression in the *mdf-1* mutant by 3 DAW (P < 0.05). When *MDF* expression was induced by estradiol, the MDF-OV samples showed significantly higher levels of *WOX5* expression (P < 0.01 by 5 and 8 DAW), showing that MDF is a positive regulator of *WOX5*. This correlated with an increase in both root length and root branch number in MDF-OV samples compared to wild type after longer periods of culture (Fig. 4, A and B), and is consistent with an inductive effect of MDF on adventitious root meristem formation.

When *MDF* expression was induced by estradiol over an 8 d regeneration time course, the MDF-OV samples showed significantly higher levels of *RAP2.7* expression (Fig. 4, E and F). In the absence of estradiol induction, there was no significant difference in *RAP2.7* expression between wild type (Col-0) and MDF-OV samples, and there was no detectable expression in the *mdf-1* mutant at any point during the experimental time course. *NAC1* expression was undetectable in leaf at 0 DAW for any genotype, but was detectable by 3 DAW, and not significantly different at this timepoint in wild type and MDF-OV, but it was significantly reduced in the *mdf-1* mutant (Fig. 3, G and H). By 5 and 8 DAW *MDF* overexpression led to a significant increase in *NAC1* expression, while *NAC1* expression was significantly reduced in the *mdf-1* mutant, compared to levels in wild type (Fig. 3, I and J). This demonstrates a role for MDF in the positive transcriptional control of both *RAP2.7* and *NAC1* following wounding.

Previous experiments has demonstrated a role for hormones other than auxin and ethylene in the promotion of AR regeneration from leaf, and notably wound-related jasmonic acid (e.g. Zhang et al., 2019, 2023), while other hormonal signals such as cytokinin, gibberellin, strigolactone and ABA are negative regulators, likely via suppression of auxin responses (Bellini et al., 2014; Jing et al. 2020). To investigate how *MDF* expression regulation might be integrated with these other signalling pathways known to be important in *de novo* AR regeneration, we analysed transcriptome data from a large number of experiments in which seedlings of Arabidopsis were treated with a wide range of hormones (data from bar.utoronto.ca; see references). The results show that *MDF* is not significantly transcriptionally regulated over a time course of between 30 min and 3 h (according to the experiment) by ACC, zeatin, IAA, ABA, methyl jasmonate, gibberellic acid (GA3), brassinolide or a number of other sterols. The lack of induction of *MDF* expression by jasmonate indicates it is regulated indepdendently of the wound response, consistent with its meristem expression in intact embryos and seedlings (Casson et al. 2009; Thompson et al. 2023). However, we have previously shown that *MDF* expression increases under abiotic stresses, to maintain seedling stem cell viability (Thompson et al. 2023), and analysis of publicly available abiotic stres-related transcriptomic data similarly show that *MDF* expression is slightly up-regulated by reactive oxygen species (ROS) after 3-6 h after treatment with 10 µM methyl viologen (Suppl. Table S3), and of interest is that ROS promote *de novo* root regeneration (Liu et al. 2023).

### The RAP2.7 transcription factor regulates expression of *WOX5* but not *YUC1* or *NAC1* in root regeneration

To investigate a possible regulatory role of the MDF target *RAP2.7*, transgenic overexpressers (*pro35S::RAP2.7*, RAP-OV) and two independent confirmed loss-of-function SALK mutants of *RAP2.7* were characterized; one representative line of each was further analysed for adventitious root initiation and growth, and for the expression of *YUC1*, *NAC1* and *WOX5* genes at 12, 19 and 26 DAW. The RAP-OV line used showed a ca. 7.4-fold increase in *RAP2.7* expression compared to wild type (Suppl. Fig. S10). At 12 DAW the mean number of roots formed was not statistically significantly different between the three genotypes. By 19 DAW the number of roots formed increased for both wild type and overexpresser, and there was no significant difference between them; however, the *rap2.7* mutant produced significantly fewer roots than wild type and showed evidence of senescence (Fig. 4B). There were significantly more roots initiated in the *RAP2.7* overexpresser, and significantly fewer in the *rap2.7* mutant, than wild type by 26 DAW. At 12 DAW, there was no statistically significant difference in root length between Col-0 and RAP-OV genotypes, but the *rap2.7* mutant produced significantly shorter roots than wildtype (Fig. 4, A and C). At both 19 and 26 DAW the *RAP2.7* overexpresser produced significantly longer roots than wild type. These results show a role for RAP2.7 in both initiation and growth of *de novo* adventitious roots and associated lateral roots in this system.

Expression levels of *YUC1*, *NAC1* and *WOX5* transcripts in the *RAP2.7* overexpresser and mutant, compared to wild type, were determined during a 7 d culture period post wounding. The results in Fig. 4E show that, for all genotypes, the level of *YUC1* expression increased during the experimental time course, but there was no statistically significant difference between mutant and overexpresser of *RAP2.7* at any time point, compared to wild type. These data show that *YUC1* expression is regulated by a pathway independent of RAP2.7. The same pattern was observed for *NAC1* expression levels in the different genotypes (Fig. 4F), again indicating that *NAC1* expression is independent of RAP2.7. However, *WOX5* expression was significantly increased in the *RAP2.7* overexpresser and decreased in the *rap2.7* mutant by 5 DAW, showing that the WOX5 pathway is positively regulated by RAP2.7.

### NAC1 regulates root regeneration independent of *YUC1*, *WOX5* or *RAP2.7*

There is previous evidence of the NAC1 transcription factor being able to promote adventitious rooting in an auxin-independent manner (Chen et al., 2016c). To understand better the mechanistic basis of the regulatory role of NAC1, transgenic overexpressers (NAC-OV) and independent confirmed loss-of-function mutants of *NAC1* were identified and one representative line of each was analysed for adventitious root formation and growth, and for the expression of *YUC1*, *WOX5* and *RAP2.7* genes during post-wounding culture of leaf. The NAC1-OV line used showed a ca. 13-fold increase in *RAP2.7* expression compared to wild type (Suppl. Fig. S11). Throughout the entire time course there was no statistically significant difference in number or length of roots formed between Col-0 and NAC-OV genotypes, but the *nac1* mutant failed to produce any adventitious roots (Fig. 5, A-C). This confirms previous reports of the requirement of NAC1 for *de novo* root formation. Expression analysis showed that, for all genotypes, the level of *YUC1*, *WOX5* and *RAP2.7* expression increased during the experimental time course, but there was no statistically significant difference in expression levels for any of the genes between mutant and overexpressers of *NAC1* compared to wild type (Fig. 5, D-F). These results show that NAC1 does not regulate *YUC1*, *WOX5* or *RAP2.7* during *de novo* root regeneration, and its role in AR formation is by a pathway independent of these genes.

**Figure 5.**
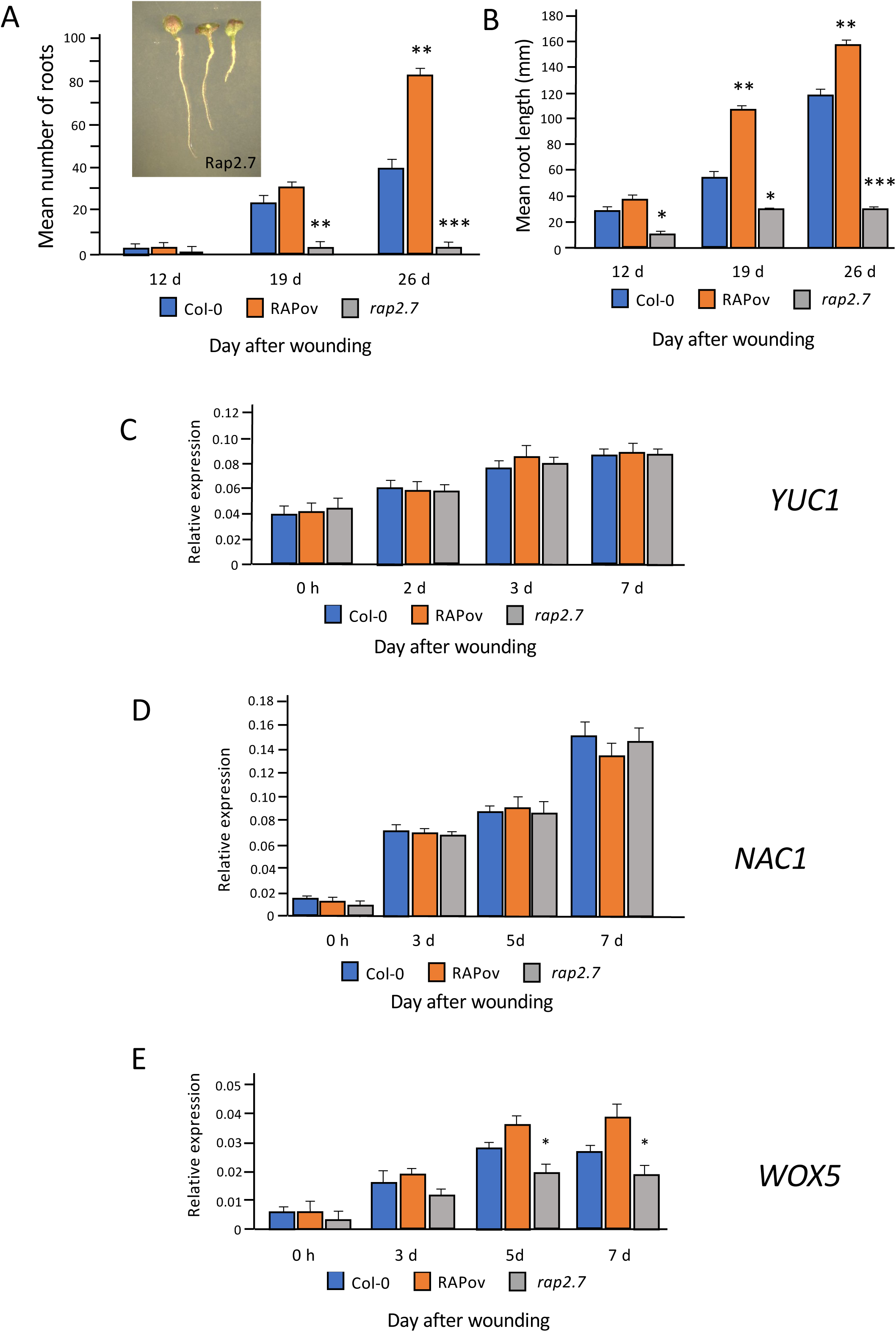
(A) Mean numbers of regenerated roots and (B) mean root lengths for wild type, *RAP2.7* overexpressers and *rap2.7* mutants at 12, 19 and 26 d after wounding on B5 medium. Inset in (A):12 day old explants of (left) *RAP2.7* overexpresser, (centre) Col-0 and (right) *rap2.7* mutant. Values are means of at least 30 samples + SEM. C-E: RT-qPCR analysis of (C) *YUC1*, (D) *NAC1* and (E) *WOX5* gene expression in wild type (Col-0), *RAP2.7* overexpresser and *rap2.7* mutant leaf during a culture time course of 0 -7 d after wounding, using *UBC* as reference gene. Values represents means and error bars are SEM (n = three biological repeats with three technical repeats). Statistical significance was determined using Student’s t-test for independent samples compared to wild type values, with P-values <0.05 (*), P <0.01 (**), P-value < 0.001 (***).

## Discussion

Plants have a remarkable developmental plasticity and capacity for regeneration. They can create new organs from non-embryonic tissue (Hartmann et al., 2010; Chen et al., 2014; Liu et al., 2014) and readily repair damage that occurs upon wounding (Xu et al., 2006; Heyman et al., 2013). Wounding plays a major role in inducing the accumulation of the hormone auxin at high concentrations close to the cut surface (Chen et al., 2016a, b). Crosstalk between auxin, cytokinin and other hormones is an important regulatory mechanism of many aspects of plant development and regeneration (El-Showk et al., 2013; Moore et al., 2015, 2024; Liu et al. 2017), including in *de novo* root formation.

There are considered to be two pathways that regulate *de novo* root regeneration from leaf explants, namely the auxin pathway and the NAC1 pathway; and past studies have shown that defects in either of these two pathways causes repression of *de novo* root development (Chen et al., 2016a-c). During *de novo* root formation cell division occurs in competent cells in the petiole and transforms these cells to root founder cells, in a process involving the expression *WOX11/12* genes (Liu et al., 2014). The root founder cells then divide to form root primordium cells, associated with expression of *WOX5/7* genes, which activate *WOX11/12* gene expression is required to activate *WOX5/7* (Hu and Xu, 2016). Finally, the root apical meristem continues to develop from the root primordium cells, leading to root emergence from the leaf explant.

MDF is a component of the plant spliceosome and is itself regulated transcriptionally in an auxin-independent manner (Casson et al., 2009; Thompson et al., 2023). It is required for the correct splicing of genes involved in meristem function, such as *RAP2.7*, *RSZ33* and *ACC1*, and for the correct level of expression of other genes, downstream of spliced targets, that themselves regulate auxin transport and auxin-mediated meristem control (Thompson et al., 2023). Lack of MDF function leads to loss of an auxin maximum in the *Arabidopsis* root due to reduced PIN protein levels and reduced of expression of other meristem genes such as *SHR* and *SCR* in an auxin-independent manner, and illustrated by reduced auxin-regulated gene (*IAA1*) expression (Suppl. Fig. S9). Previously we have shown that *MDF* overexpression could induce ectopic shoot meristems (Casson et al., 2009). In the current paper we show that MDF is required for *de novo* root formation and the correct quantitative expression of *RAP2.7*, *WOX5* and *NAC1* (Fig. 4). *MDF* overexpression increases both the expression of these genes and also the number of *de novo* roots initiated compared to wild type. RAP2.7 itself can induce both *WOX5* expression by 5 DAW and root initiation and growth by 12-19 DAW, but has no effect on the expression of either *YUC1* or *NAC1* (Fig. 5). While *NAC1* is essential for *de novo* root regeneration, it is not required for *YUC1*, *WOX5* or *RAP2.7* expression, and is not a positive regulator of these genes (Fig. 6).

**Figure 6.**
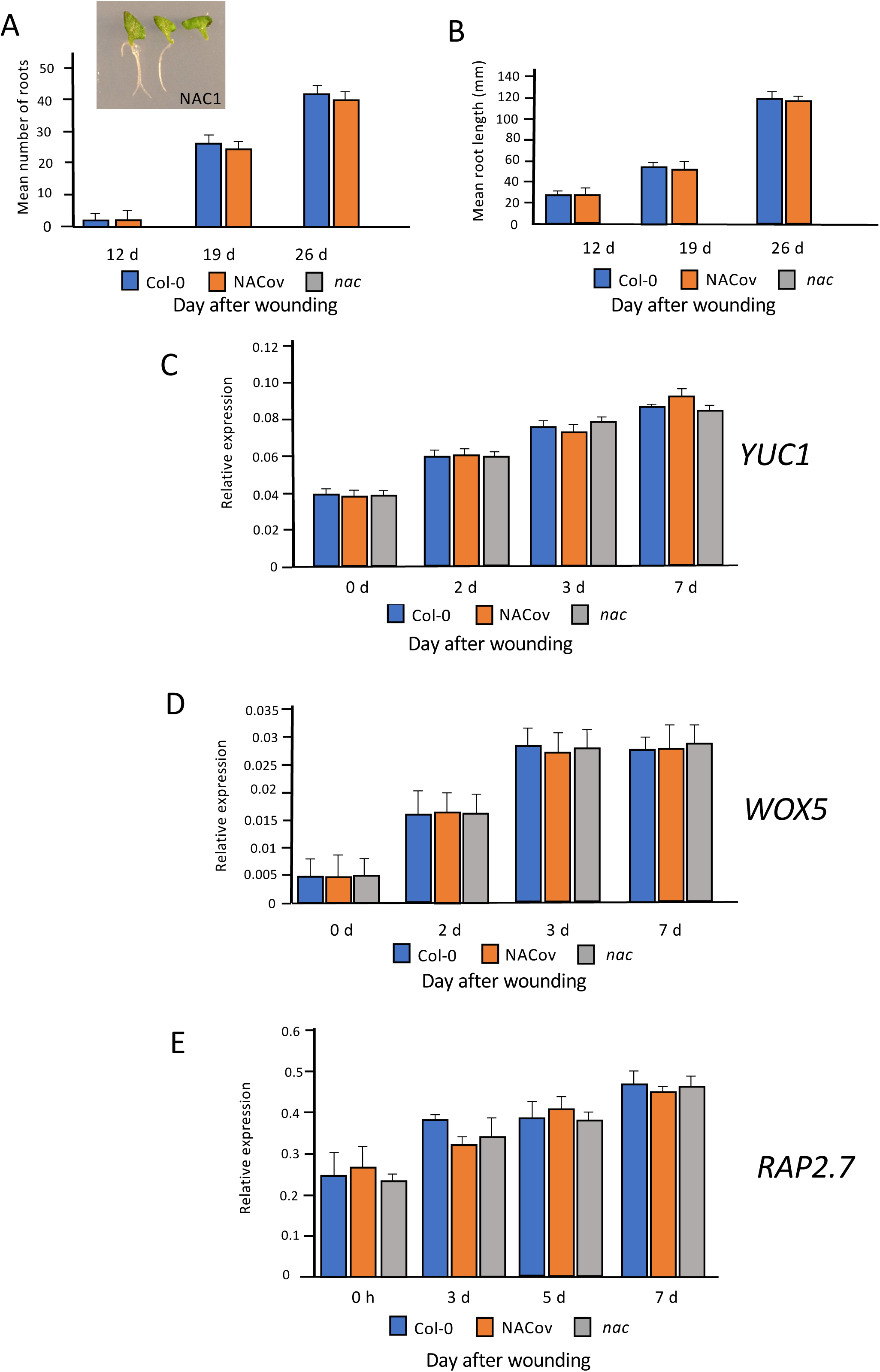
(A) Mean numbers of regenerated roots and (B) mean root lengths for wild type, *NAC1* overexpressers and *nac1* mutants at 12, 19 and 26 d after wounding on B5 medium. Inset in (A):12 day old explants of (left) Col-0, (centre) *NAC1* overexpresserand (right) *nac1* mutant. Values are means of at least 30 samples + SEM. C-E: RT-qPCR analysis of (C) *YUC1*, (D) *WOX5* and (E) *RAP2.7* gene expression in wild type (Col-0), *NAC1* overexpresser and *nac1* mutant leaf during a culture time course of 0 -7 d after wounding, using *UBC* as reference gene. Values represents means and error bars are SEM (n = three biological repeats with three technical repeats). Statistical significance was determined using Student’s t-test for independent samples compared to wild type values; no significant difference between samples was found.

We show that the NAC1 pathway is not dependent on either auxin or ethylene signalling, whereby *NAC1* expression was unaffected during *de novo* root development in mutants in either auxin signalling (*axr1*, *axr3*) or ethylene signalling (*ein2*, *pls*) (Fig. 3). The *pls* mutant is an ethylene hypersignalling mutant that also has low auxin concentrations in the root (Chilley et al., 2006), consistent with its observed low expression of *YUC1* and *4*, associated with defective *de novo* root initiation and growth (Figs. 1, 3). The *EBS::GUS* ethylene reporter was expressed in the petiole but at lower levels in the emerging root in the wild type background (Suppl. Fig. S6), consistent with low ethylene responses being associated with efficient root elongation. Root formation from leaf of the ethylene-insensitive *ein2* genotype was similar to wild type, consistent with previous results suggesting that the *ein3* mutant, also ethylene-insensitive, was able to form more *de novo* roots than wild type, and it was proposed that the ethylene hormone is a negative regulator of *de novo* root formation (Li et al., 2021). While there is good evidence that ethylene inhibits root growth (e.g. Casson et al., 2002; Lewis et al., 2011), the results here for *ein2* are not completely in agreement with the data for *ein3*, as *ein2* does not regenerate more or longer roots than wild type, showing that *de novo* root formation is not promoted by ethylene insensitivity; and indeed there may be fewer root branches from the *ein2* leaf than for wild type at 26 d of culture (Fig. 1). This suggests some ethylene sensitivity/signalling activity is required for root initiation over a prolonged culture period, possibly through crosstalk with other hormones (Moore et al., 2024).

The *ein2* genotype has no effect on *YUC1* and *YUC4* expression during leaf explant culture (Fig. 3). Previous results showed that the ethylene-stabilized transcription factor EIN3 directly represses the expression of both *WOX11* and *WOX5*, which are key cell fate-determining genes, providing a possible mechanism for ethylene-mediated repression of *de novo* root formation (Li et al., 2021). Similarly, ERF115 (ETHYLENE RESPONSE FACTOR 115) represses root regeneration via effects on jasmonate and cytokinin signalling (Lakehal et al., 2020). Furthermore, ethylene signalling can reduce auxin accumulation to cause reduced root regeneration (Fig. 1) (Li et al., 2021). This possibility is consistent with our results which show *WOX5, YUC1* and *WUC4* expression is reduced in the ethylene hypersignalling and low auxin *pls* mutant; i.e. ethylene responses are repressed to allow root initiation (Fig. 3). Interestingly, a short treatment of excised leaves of Arabidopsis with either ethylene (Liu et al., 2022) or the ethylene precuror ACC can promote *de novo* AR initiation in Petunia (Druege et al., 2014), but ethylene inhibits later regeneration events (such as root growth and lateral root formation), indicating that a tight control over ethylene signalling is required for regeneration. Past studies have shown that there is crosstalk between ethylene and auxin in the root, whereby ethylene induces two Trp biosynthetic genes, *WEI2/ASA1/TIR7* and *WEI7/ASB1*, which leads to increased auxin concentrations in the root tip and elongation zone, which is inhibitory to growth (Stepanova et al., 2007). It can be proposed that excess ethylene is inhibitory to the root regeneration process, but ethylene insensitivity has little adverse effect, and signalling needs to be maintained to allow efficient adventitious root formation to occur, likely in part at least through the requirement for ethylene for correct auxin and cytokinin patterning (Moore et al., 2024).

These results suggest a regulatory network in which MDF regulates both auxin-dependent (via *RAP2.7* and *WOX5*, as well as e.g. *PIN* and *PLT* genes) and auxin-independent (via *NAC1*, as well as e.g. *SHR* and *SHR;* Casson et al., 2009, Thompson et al., 2023) pathways of *de novo* root regeneration (Fig. 7). *AXR1* is essential for auxin responses and auxin biosynthesis via YUC1 and YUC4, required for root regeneration (Fig. 3), as a central component of the auxin response pathway (Kubalova et al., 2024). Interestingly, AXR3 appears to play a less critical role, as the gain-of-function *axr3-1* mutant shows higher frequency root initiation than the *axr1* mutant, with less severe negative effects on the expression of *WOX5*, *YUC1* and *YUC4* auxin pathway genes than *axr1* (Fig. 3). ROS may have some effect in positively regulating *MDF* expression (Suppl. Table S2), associated with the known promotive effect of ROS on *de novo* root regeneration (Liu et al. 2022). Further studies are needed to understand how MDF is itself regulated during development, to control diverse downstream pathways.

**Figure 7.**
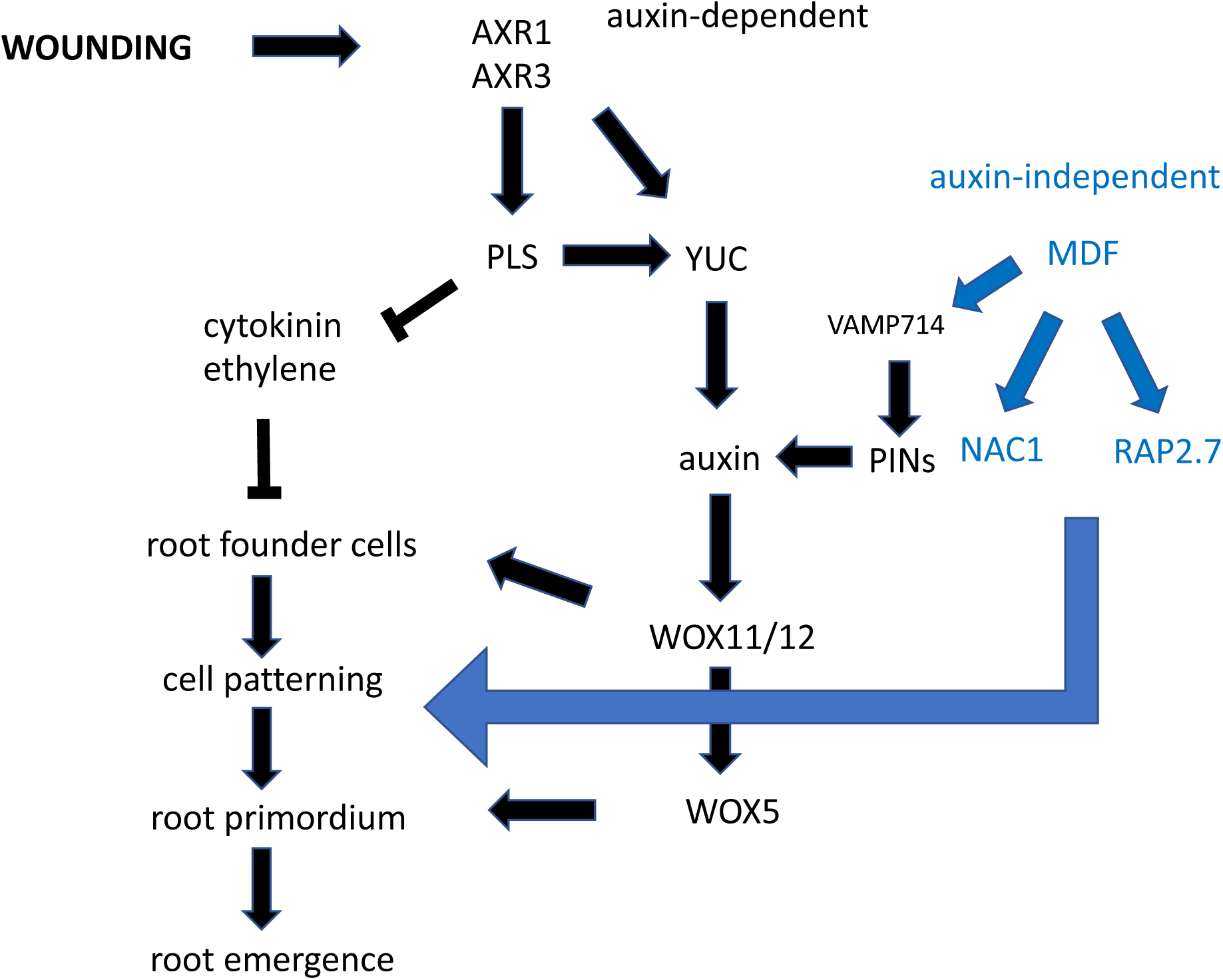
Summary network model for data integrated with known information on gene-hormone interactions. Wounding induces the AXR/auxin-dependent and MDF/auxin-independent pathways. These interact at the interface of PINs, which require MDF activity and regulate auxin transport produced by the YUC pathway. Auxin activates the *WOX* genes and MDF activates the independently regulated NAC1 and RAP2.7 pathways. *PLS* is positively regulated by auxin and is a negative regulator of both ethylene and cytokinin, to suppress the repressive effects of these hormones on root development.

## Materials and Methods

### Plant material

Wild type (WT) *Arabidopsis thaliana* Columbia (Col-0) seeds were obtained from lab stocks, Department of Biosciences, Durham University. All mutant lines were also obtained from lab stocks - *mdf-1*, *axr1*, *axr3* and *ein2* (T-DNA Salk mutant lines). The *pls* mutant is a promoter trap T-DNA insertion line (Casson et al. 2002). GUS reporter lines and Salk mutant lines were obtained from the NASC website (http://arabidopsis.info).

Seedlings were grown *in vitro* on half strength Murashige and Skoog medium (1/2MS10) (Murashige & Skoog, 1962) containing 2.2 g/l half strength MS medium (Sigma M5519) with 10g/l of sucrose, adjusted to pH 5.7, and 8 g/l agar. For leaf culture, B5 medium (Sigma G5893) was used (3.2 g/l of B5 medium, adjusted to pH 5.7, solidified with 8 g/l agar). All the media were sterilized in the autoclave at 121°C, 1.1 bar for 20 min.

For regeneration experiments, seedlings or isolated leaves were cultured on solidified media and incubated in the growth room or growth cabinets under 16 h light: 8 h dark at 22°C (c. 3000 lux). Seeds were germinated and grown for 12 d on sterile square Petri dishes (size 10 x 10 cm) containing 50 ml of solid half MS medium. Then, after 12 d the first two leaves were removed with sterilized mini scissors and the leaves were transferred to B5 medium using sterilized forceps. For assessment of root regeneration, *de novo* adventitious roots formed typically up to 12 days after wounding, and may continue to grow and develop lateral roots thereafter; the counts up to 12 days after wounding therefore represent *de novo* adventitious root numbers, but total root numbers and adventitious primary root lengths were also determined. Root lengths were measured on an Epson Expression 1680Pro flatbed scanner (Epson, UK) set at a resolution of 800 dpi. Measurements were quantified using ImageJ software (Schneider et al., 2012) with the ‘smartroot’ plugin (Shahzad et al., 2018).

For gene expression comparisons at different times, the leaves were collected on 0 h, 3 d, 5 d and 7 d (in the signalling experiment the leaves were collected on 0 h, 24 h, 48 h, 3 d, 5 d, 7 d,) and stored at -80°C prior to RNA extraction.

### Gene cloning

An inducible *MDF* overexpressor transgenic line was designed for comparative studies with the loss-of-function *mdf-1* mutant and constructed using Gateway cloning (Invitrogen). All the primers were designed by SnapGene software and sequences are available in Suppl. Table S3. The pDONR207 vector was used for all the BP reactions, according to the manufacturer’s instructions.

An *MDF* entry clone from lab stocks, containing the coding sequence of the *MDF* gene (Thompson et al., 2023), was used to produce the expression clone to overexpress *MDF* under the control of an estradiol inducible promoter *XVE-35S* in the vector pMDC7. This was expected to ensure the overexpression of the gene. The purified entry clone was used for LR reaction, cloned into the pMDC7 destination vector, and introduced into chemically competent DH5a cells according to the Gateway instructions. Positive colonies were confirmed by PCR. The correct *XVE-35S::MDF* expression clone was introduced into *Agrobacterium tumefaciens* strain GV3101 by the freeze-thaw method, and positive colonies were identified by their ability to grow on LB-agar plates with rifampicin, gentamycin, and kanamycin antibiotics. Arabidopsis was transformed by the method of Clough and Bent (1998).

Plants expressing *MDF*, which showed 3:1 segregation in the T2 generation (indicative of transgene insertion at a single locus), were screened for homozygous identification. Individual T2 plants were grown until seed development. Then, seeds were sown on hygromycin plates. Lines that showed 100% resistance to the antibiotic and able to overexpress *MDF* were identified as homozygous T3 seedlings. The line *XVE-35S::MDF-H3* was selected as the overexpressor line, from now on will refer to this line as MDF-H3. RT-qPCR was performed to confirm the overexpression after induction with estradiol.

*RAP2.7* (AT2G28550.1) and *NAC1* (AT1G56010.1) transgenic overexpressers were also made. All the primers and entry cloning sequences for Gateway pDONR207 were designed using the Snapgene programme. In addition, different forward primers at different sites in the *RAP2* and *NAC* genes were designed to confirm entry cloning following PCR cloning and sequencing. The list of primers is presented in Suppl. Table S3. Purified entry clones were used for LR reactions, cloned into the pMDC7 destination vector, and introduced into chemically competent DH5a cells according to the Gateway instructions. Positive colonies were confirmed by PCR and sequencing. The correct clones was introduced into *Agrobacterium tumefaciens* strain GV3101 by the freeze-thaw method, and positive colonies were identified by their ability to grow on LB-agar plates with rifampicin, gentamycin, and kanamycin antibiotics. Arabidopsis was transformed by the method of Clough and Bent (1998).

### Estradiol induction

β-estradiol was used for induction of *XVE-35S::MDF* gene expression according to Zuo et al. (2000). For leaf induction treatments, the hormone was a final concentration of 5 µM and was sprayed onto the leaf cultured on B5 medium. Then the leaf was collected after 24 h induction. For 0 h leaf the seedling on 11 d culture on half MS medium was sprayed with hormone; and after 24 h the leaf was collected after 12 d culture on half MS medium.

### Quantitative RT-PCR (RT-qPCR)

RNA extraction and cDNA synthesis for RT-qPCR was carried out using wild type and mutant or transgenic seedlings essentially as described previously (Rowe et al., 2016), using three biological and three technical replicates. qPCRBIO SyGreen Mix Lo-ROX kit (PCR BIosystems) was used for qPCR analysis, and the reaction mixture and the programme conditions were set according to the supplier’s instructions and run on a Rotor-Gene Q Machine (QIAGEN). *UBIQUITIN C* (*UBC*)*, UBIQUITIN10* (*UBQ10*) or *ACTIN2* (*ACT2*) were used as reference genes. Expression analysis was conducted using the Rotorgene Q Series software v1.7. Relative normalised levels of transcript of each gene were calculated relative to the reference gene and analysed by comparative quantification using an assumption-free, linear regression analysis approach (Ramakers et al., 2003). Primer sequences are listed in Suppl. Table S3.

### Bioinformatics

Gene expression data for the *MDF* gene regulation in response to hormones and abiotic stresses (Suppl. Tables S1 and S2) were obtained from https://bar.utoronto.ca/efp/cgi-bin/efpWeb.cgi?dataSource=Hormone&mode=Absolute&primaryGene=At5g16780&secondaryGene=At3g27340&override=&threshold=604.48&modeMask_low=None&modeMask_stddev=None). *MDF* and *RAP2.7* expression during *de novo* root regeneration were obtained from RNA-seq data published by Liu et al. (2023).

### Histochemistry and microscopy

GUS histochemistry was carried out as described previously (Topping and Lindsey, 1997). Leaf samples were examined using a Zeiss Axioskop compound light microscopy (Carl Zeiss, Cambridge, UK), equipped with a QImaging Retiga-2000r camera (Photometrics, Marlow, UK) using a x10 0bjective.

### Statistical analysis

Statistical significance was determined using either Student’s t-test for independent samples compared to wild type values, with P-values <0.05 (*), P <0.01 (**), P-value < 0.001 (***); or one-way ANOVAs and correlation analysis were performed using SPSS 26.0 software, and Duncan ’s new multiple range method was used for significance test (P < 0.05 ). Excel 2018 software was used to tabulate basic data, and Origin 2021 software was used to draw the single factor histogram.

## Acknowledgements and funding

We are grateful for funding from the Biotechnology and Biological Sciences Research Council (grants BB/S000305/1 and BB/M011186/1 to K.L. and a studentship to JA); from the Saudi Arabian government for a studentship to FM; and from Consejo Nacional de Ciencia y Tecnología (CONACYT) and the Secretaría de Energía de México for a studentship to R.M. We thank the Durham Genomics facility, Department of Biosciences, for DNA sequencing.

## Author contributions

KL, JFT: devised the project; FA, RM, HZ: carried out the experimental work and prepared the figures; KL, JFT, JA: supervised the work; KL: produced the first draft of the manuscript; all authors contributed text and edited the manuscript.

## Supplementary material

Figure S1. Expression of the auxin reporter *DR5::GUS* during *Arabidopsis* adventitious root regeneration.

Figure S2. *PIN1* gene expression in *Arabidopsis* leaf and regenerating root.

Figure S3. *PIN3* gene expression in *Arabidopsis* leaf and regenerating root.

Figure S4. *PIN7* gene expression in *Arabidopsis* leaf and regenerating root.

Figure S5. *VAMP714::GUS* gene expression in *Arabidopsis* leaf and regenerating root.

Figure S6. *EBS::GUS* gene expression in *Arabidopsis* leaf and regenerating root.

Figure S7. Expression of *MDF* and *RAP2.7* genes as determined by RNA-seq over a 5 day leaf culture preiod for adventitious root regeneration.

Figure S8. *MDF* inducible gene expression.

Figure S9. Relative expression of *IAA1* in *mdf* mutant and WT (Col-0) analysed using qRT-PCR in whole leaf at 0d, 2d, 4d, 6d, 8d using *ACTIN2* as the reference gene.

Figure S10. *RAP2.7* expression levels in *35S::RAP2.7* transgenics.

Figure S11. *NAC1* expression levels in *35S:NAC1* transgenics.

**Table S1.** Gene expression data for the *MDF* gene regulation in response to hormones.

**Table S2.** Gene expression data for the *MDF* gene regulation in response to abiotic stresses.

**Table S3.** PCR primers.

