## Supplementary Figures for "MDF regulates a network of auxin-dependent and -independent pathways of adventitious root regeneration in *Arabidopsis*"

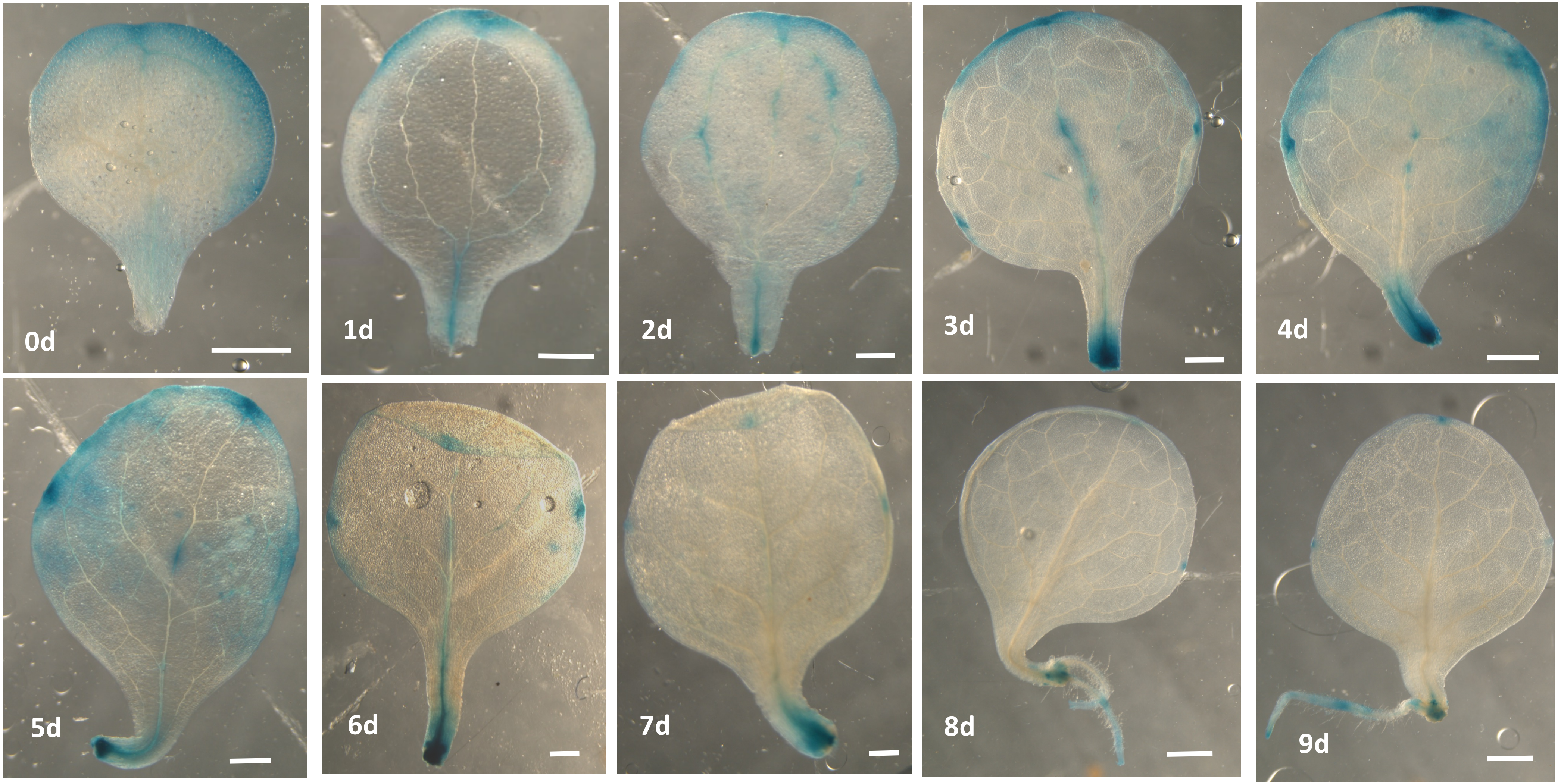


**Fig. S1. Expression of the auxin reporter *DR5::GUS* during *Arabidopsis* adventitious root regeneration.** Scale bars =500 um.

### Fig. S2. *PIN1* expression in *Arabidopsis* leaf and regenerating root.

A

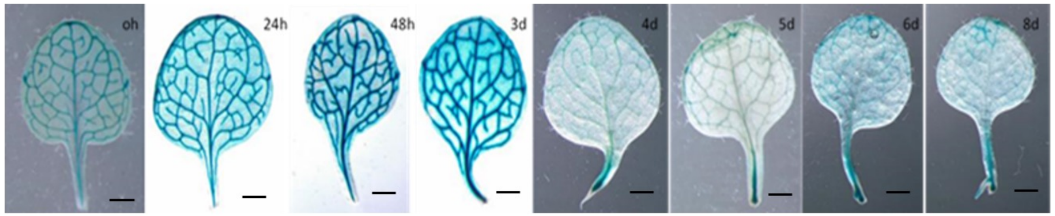


B

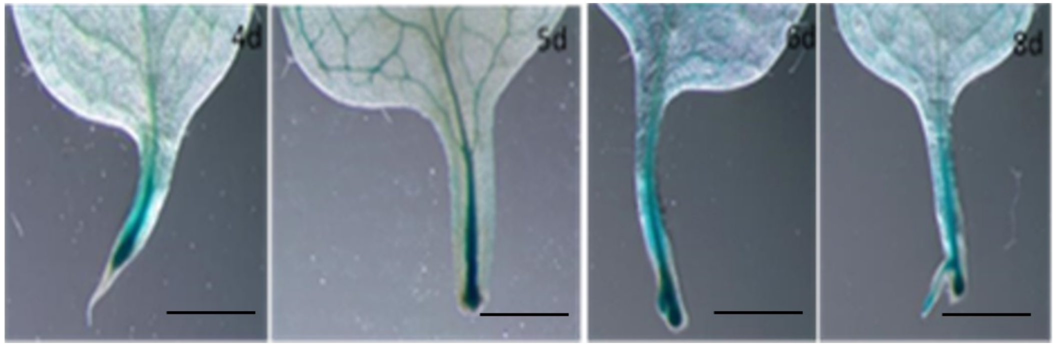


(A) *PIN1::GUS* expression in leaf of Arabidopsis when cultured from 0 h to 8 d, with scale bars = 1 mm. (B) Magnification of images in A, showing PIN1::GUS expression in petiole of leaf of Arabidopsis from 4 d to 8 d, with scale bars = 0.5 mm.

C

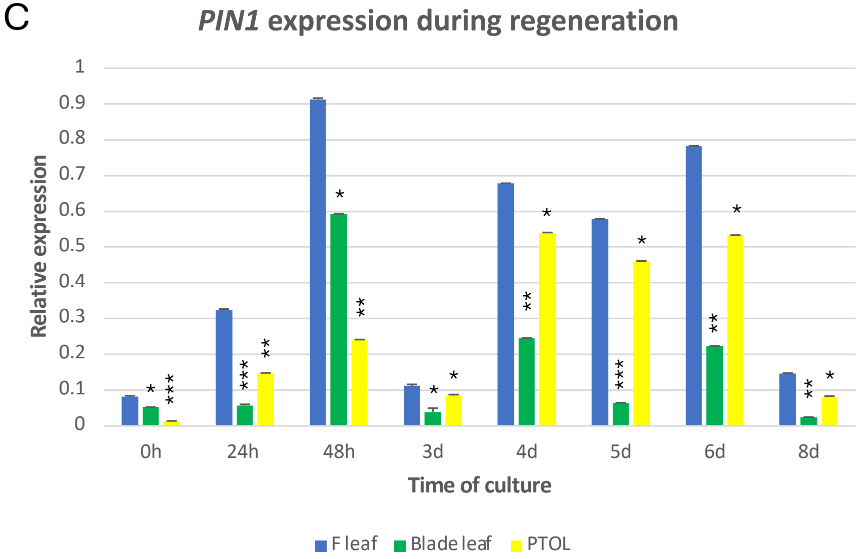


C. Relative expression of the *PIN1* gene analysed using qRT-PCR in whole leaf (blue bars, F leaf), leaf blade (green bars) and petiole (yellow bars, PTOL) from 0 h to 8 d by using the *UBC* reference gene. Values represents means and error bars are SEM (n = three biological repeats with three technical repeats). Statistical significance was determined using Student's t-test for independent samples compared to wild type values.

**Fig. S3. *PIN3* expression in *Arabidopsis* leaf and regenerating root.**

A

B

C

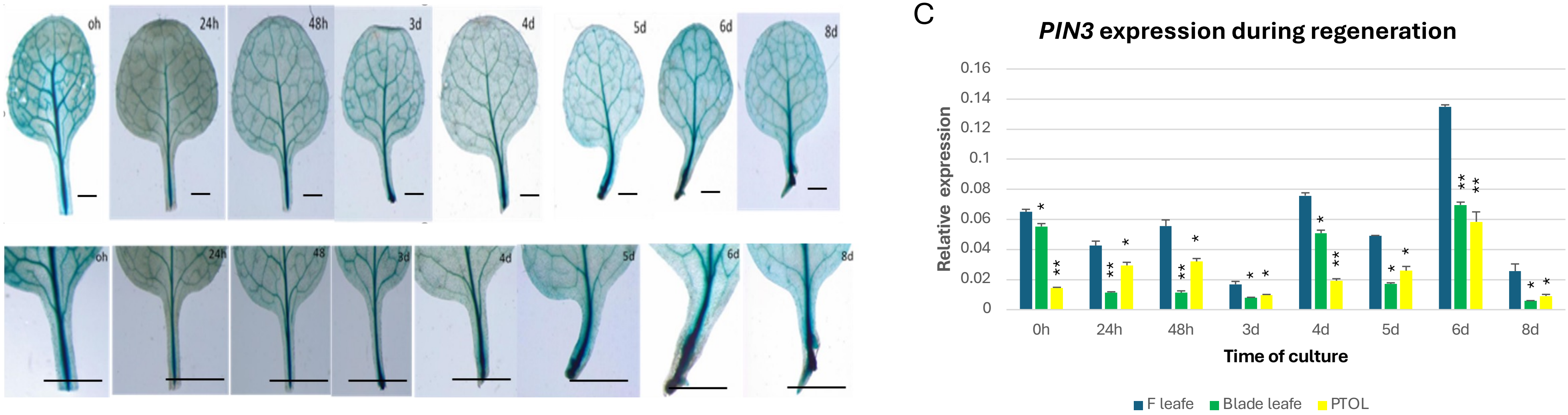


(A) *PIN3::GUS* expression in leaf of Arabidopsis when cultured from 0 h to 8 d, with scale bars = 1mm. (B) Magnification of images in A, showing *PIN3::GUS* expression in petiole of leaf of Arabidopsis from 0 h to 8 d, with scale bars = 0.5 mm.

Relative expression of the *PIN3* gene analysed using qRT-PCR in whole leaf (blue bars), leaf blade (green bars) and petiole (yellow bars, PTOL) from 0 h to 8 d using the *UBC* reference gene. Values represent means and error bars are SEM (n = three biological repeats with three technical repeats). Statistical significance was determined using Student's t-test for independent samples compared to wild type values.

**S4. *PIN7::GUS* expression in *Arabidopsis* leaf and regenerating root.**

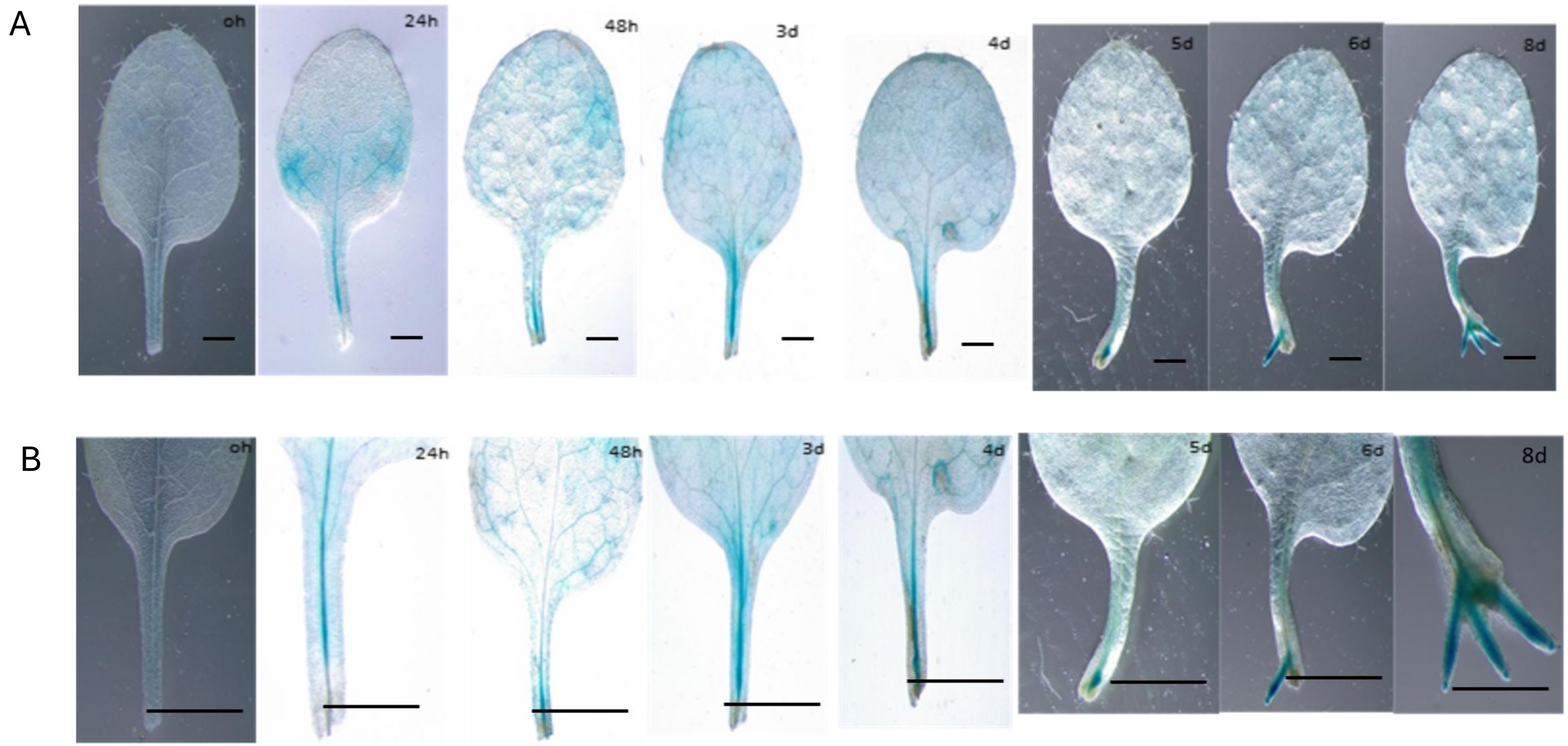


(A) *PIN7::GUS* expression in leaf of Arabidopsis when cultured from 0 h to 8 d, with scale bars = 1 mm. (B) Magnification of images in A, showing *PIN7::GUS* expression in petiole of leaf of Arabidopsis from 0 h to 8 d, with scale bars = 0.5 mm.

**Fig. S5. *VAMP714::GUS* expression in *Arabidopsis* leaf and regenerating root.**

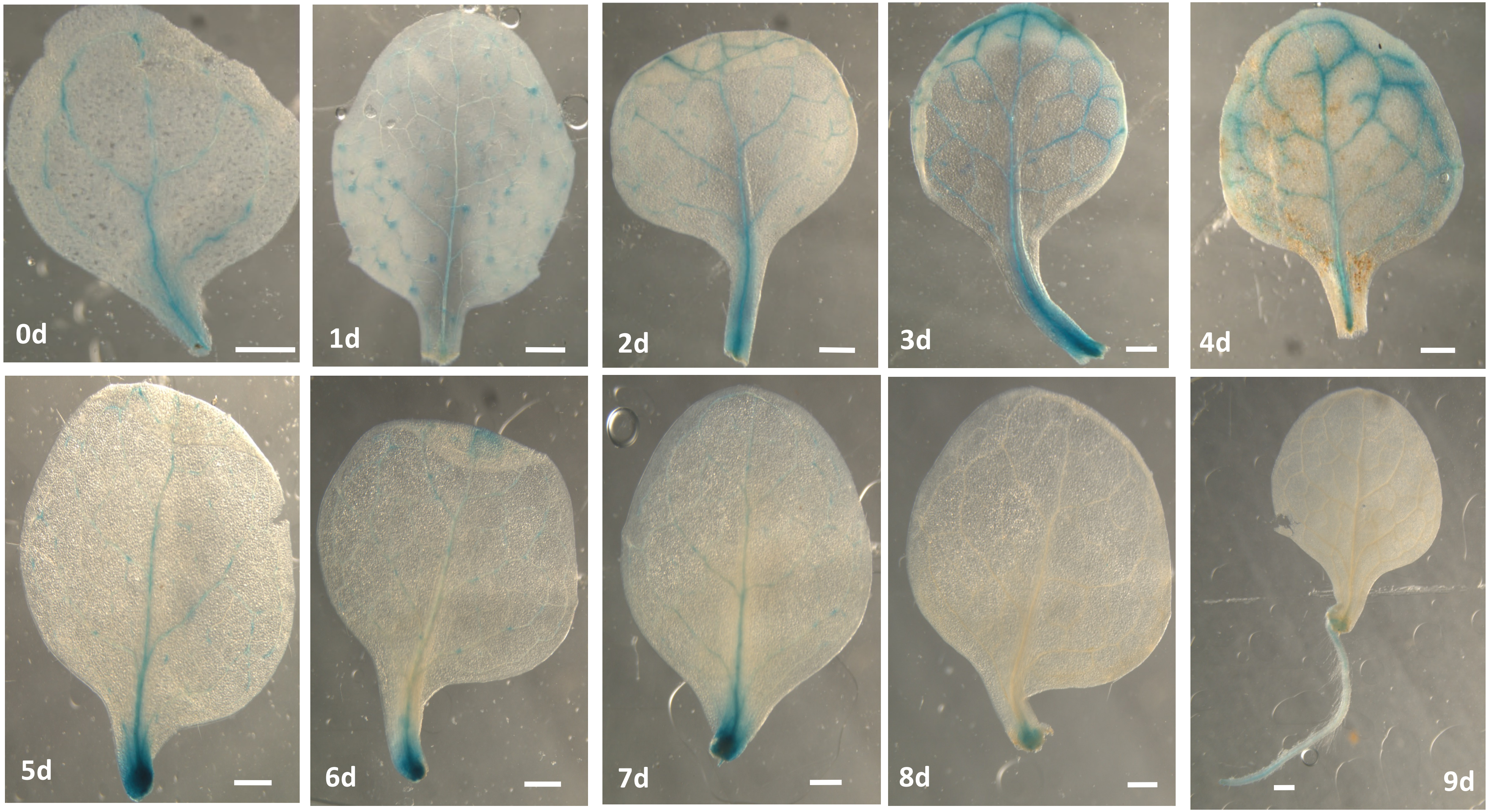


scale bars = 500 um

**Fig. S6. *EBS::GUS* expression in *Arabidopsis* leaf and regenerating root.**

A

B

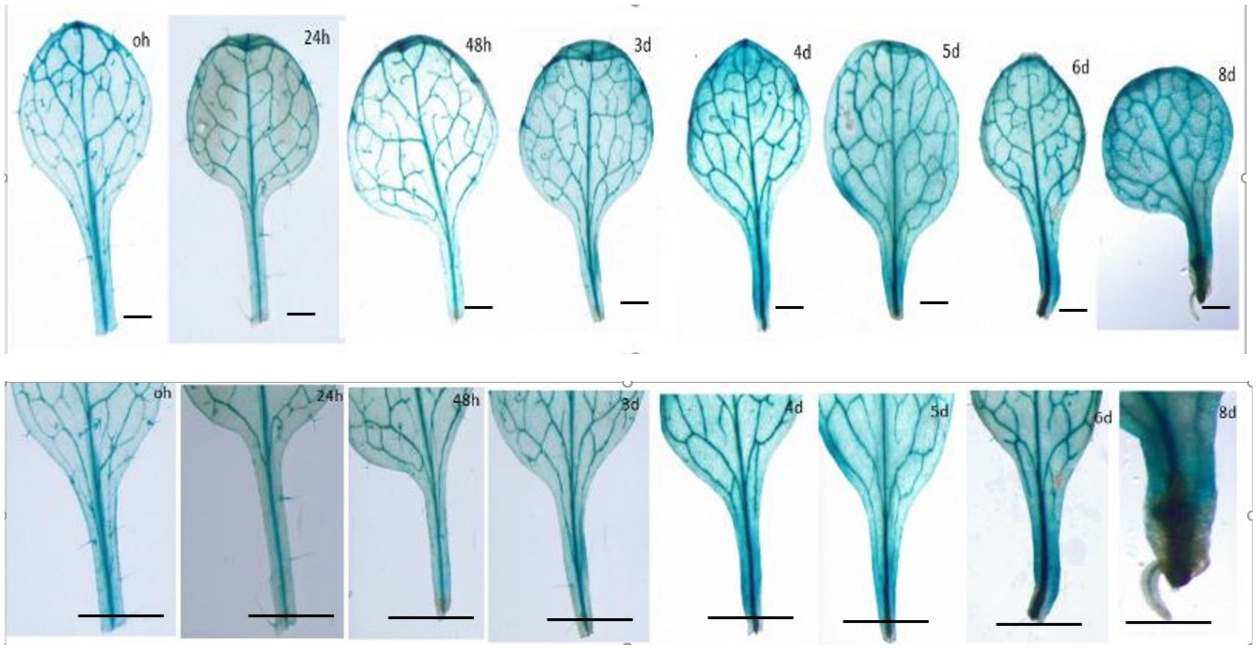


Fig. S6. A) *EBS::GUS* expression in leaf of Arabidopsis when cultured from 0 h to 8 d, with scale bars = 1 mm. (B) Magnification of images in A, showing *EBS::GUS* expression in petiole of leaf of Arabidopsis from 0 h to 8 d. Scale bars = 0.5 mm.

*MDF* (AT5G16780) expression in *Arabidopsis thaliana* detached leaves cultured on sucrose-free B5 medium (mock, without NPA) at 22°C

A

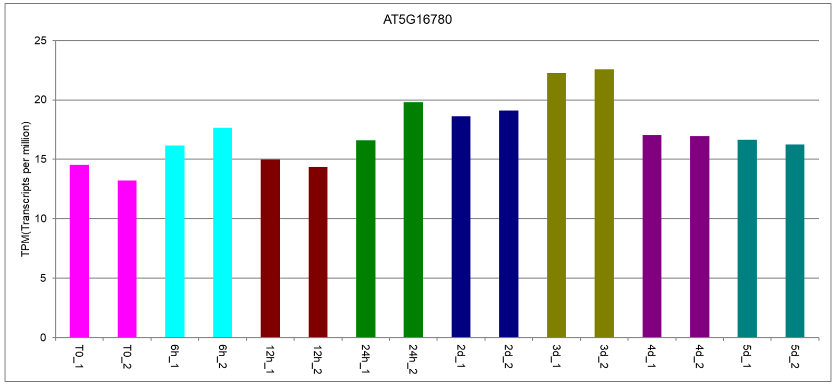


*MDF* (AT5G16780) expression in *Arabidopsis thaliana* detached leaves cultured on sucrose-free B5 medium with NPA at 22°C

B

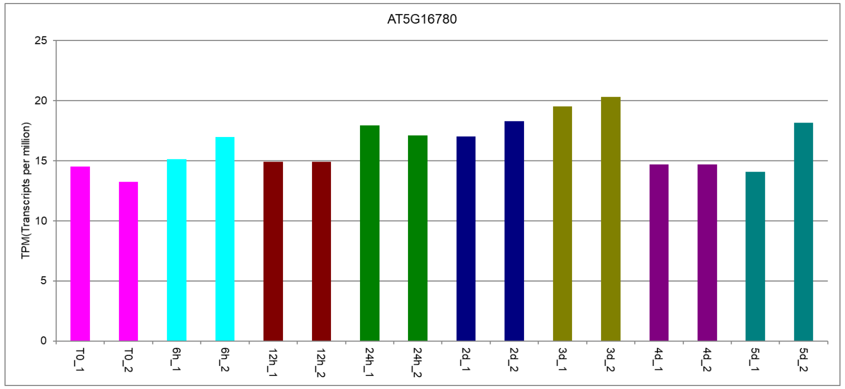


*RAP2.7* (AT2G28550) expression in *Arabidopsis thaliana* detached leaves cultured on sucrose-free B5 medium (mock, without NPA) at 22°C

C

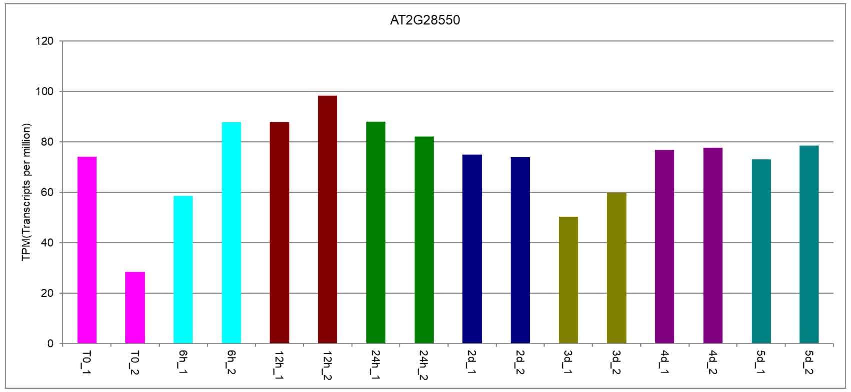


*RAP2.7* (AT2G28550) expression in *Arabidopsis thaliana* detached leaves cultured on sucrose-free B5 medium with NPA at 22°C

D

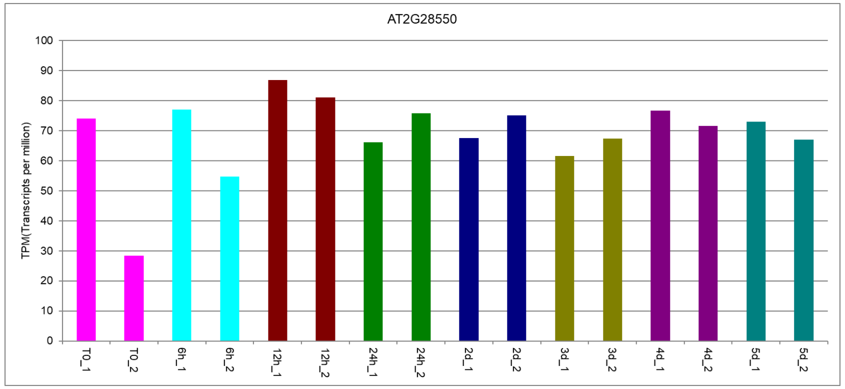


**Fig. S7**

**Fig. S7.** Expression of *MDF* and *RAP2.7* genes as determined by RNA-seq over a 5 day leaf culture period for adventitious root regeneration. Bars of the same colour represent biological replicates, at 0 h (T0, pink bars), 6 h (pale blue bars) 12 h (brown bars), 24 h (green bars), 2 d (dark blue bars), 3 d (olive green bars), 4 d (purple bars) and 5 d (teal bars).
Data from Liu et al. (2022) Plant Comms. 3,100306.

#### Fig. S8. *MDF* inducible expression

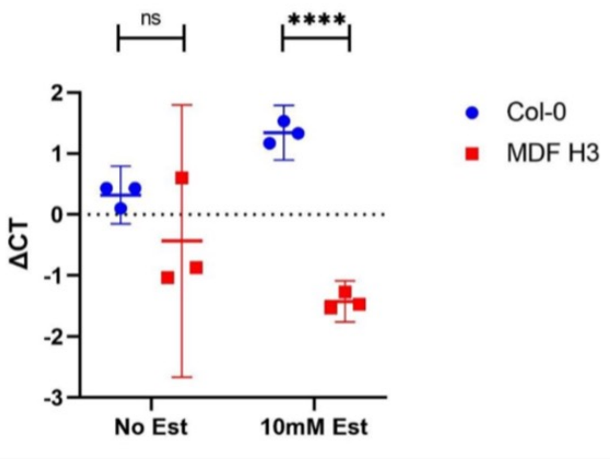


qRT-PCR analysis of *MDF* gene expression in wild type and MDF-OV. Using UBC reference gene was quantified without treatment and after treatment with 10 mM estradiol. t-test with no treatment (p = 0.28, df: 2.180); 10 mM estradiol (****p= 0.000051, df 3.720). Data from Thompson et al. (2023).

**Delta-delta Ct calculation: No estradiol induction**

| **Sample** | **Gene of interest (*MDF*)** | **Housekeeping gene (*UBC*)** | | | **Relative fold change** |
|---|---|---|---|---|---|
| | Average Ct | Average CT | ΔCt | ΔΔCt | 2^-(ΔΔCt) |
| **Col-0 A** | 20.73 | 20.30 | 0.43 | 0.11 | 0.92 |
| **Col-0 B** | 21.60 | 21.17 | 0.43 | 0.11 | 0.92 |
| **Col-0 C** | 21.47 | 21.37 | 0.10 | -0.22 | 1.16 |
| **MDF-H3 A** | 22.10 | 21.50 | 0.6 | 0.28 | 0.82 |
| **MDF-H3 B** | 20.53 | 21.40 | -0.8667 | -1.19 | 2.28 |
| **MDF-H3 C** | 20.93 | 21.97 | -1.0333 | -1.35 | 2.56 |

**Delta-delta Ct calculation: Induction with 10 mM estradiol**

| **Sample** | **Gene of interest (*MDF*)** | **Housekeeping gene (*UBC*)** | | | **Relative fold change** |
|---|---|---|---|---|---|
| | Average Ct | Average CT | ΔCt | ΔΔCt | 2^-(ΔΔCt) |
| **Col-0 A** | 24.2 | 22.67 | 1.53 | 0.19 | 0.87 |
| **Col-0 B** | 23.93 | 22.6 | 1.333 | -0.01 | 1 |
| **Col-0 C** | 23.53 | 22.37 | 1.1667 | -0.17 | 1.13 |
| **MDF-H3 A** | 21.43 | 22.7 | -1.2667 | -2.61 | 6.09 |
| **MDF-H3 B** | 21.27 | 22.73 | -1.4667 | -2.81 | 7 |
| **MDF-H3 C** | 21.1 | 22.63 | -1.533 | -2.87 | 7.33 |

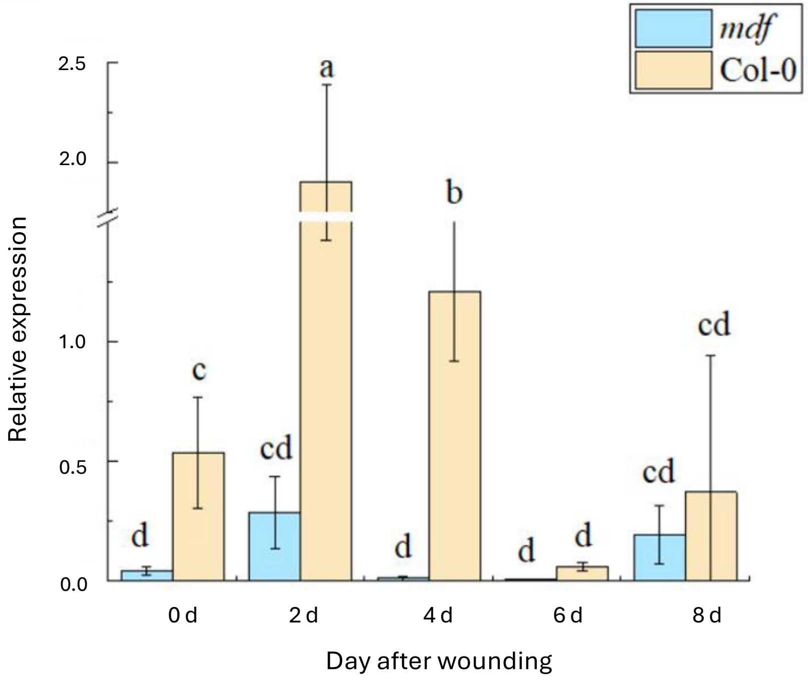


**Fig. S9. Relative expression of *IAA1* in *mdf* mutant and WT (Col-0) analysed using qRT-PCR in whole leaf at 0d, 2d, 4d, 6d, 8d using *ACTIN2* as the reference gene.** Values represent means and error bars are SEM (n = three biological repeats with three technical repeats). One-way ANOVA and correlation analysis was performed using SPSS 26.0 software, and Duncan 's new multiple range method was used for significance test ( $p< 0.05$ ).

**Fig. S10. *RAP2.7* expression levels in *35S::RAP2.7* transgenics.** Bars represent SE of mean, n = 3.

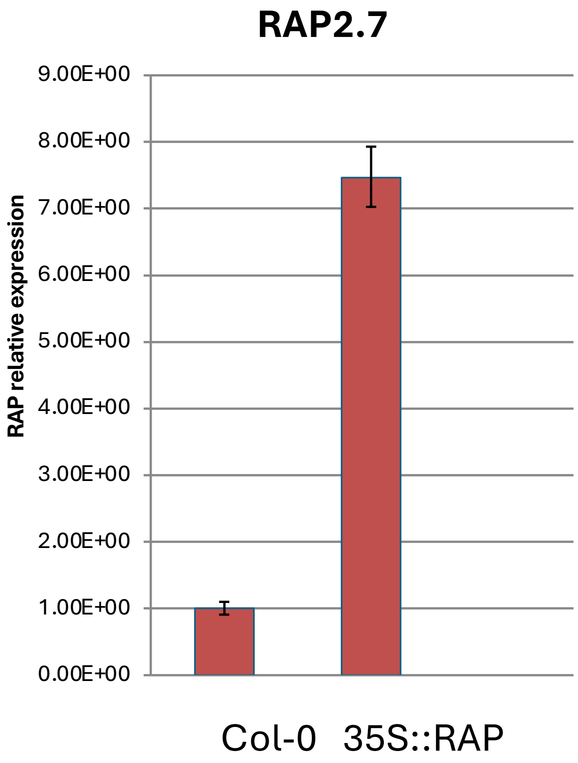

**Fig. S11. *NAC1* expression levels in *35S:NAC1* transgenics.** Bars represent SE of mean, n = 3.

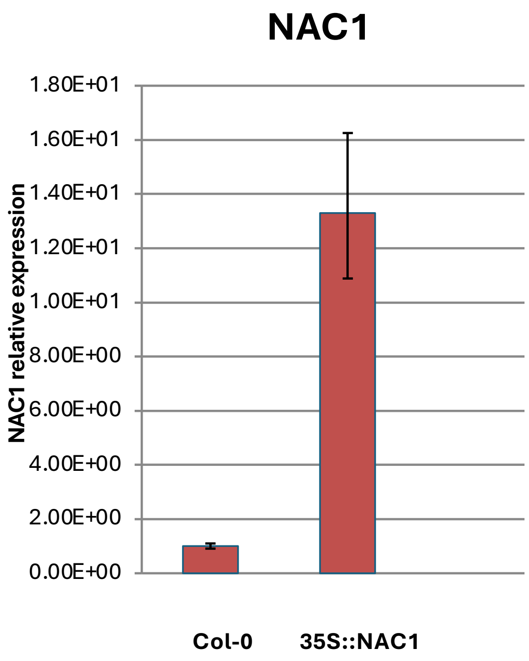
