## Supplementary material for "MDF regulates a network of auxin-dependent and -independent pathways of adventitious root regeneration in *Arabidopsis*": Table S1

| Group # | Tissue | Expression Level | Standard Deviation | Samples | Links |
|---|---|---|---|---|---|
| 1 | Control at 30 Minutes | 72.83 | 14.07 | RIKEN-GODA1AA, RIKEN-GODA1BB | To the Experiment |
| 1 | ACC Treated at 30 Minutes | 65.56 | 3.77 | RIKEN-GODA7A, RIKEN-GODA7B | To the Experiment |
| 2 | Control at 1 Hour | 80.2 | 1.99 | RIKEN-GODA9AA, RIKEN-GODA9BA | To the Experiment |
| 2 | ACC Treated at 1 Hour | 63.23 | 2.64 | RIKEN-GODA15A, RIKEN-GODA15B | To the Experiment |
| 3 | Control at 3 Hours | 65.45 | 7.44 | RIKEN-GODA17AA, RIKEN-GODA17BA | To the Experiment |
| 3 | ACC Treated at 3 Hours | 69.01 | 1.32 | RIKEN-GODA23A, RIKEN-GODA23B | To the Experiment |
| 4 | Control at 30 Minutes | 72.83 | 14.07 | RIKEN-GODA1AA, RIKEN-GODA1BB | To the Experiment |
| 4 | Zeatin Treated at 30 Minutes | 71.67 | 13.48 | RIKEN-GODA3A, RIKEN-GODA3B | To the Experiment |
| 5 | Control at 1 Hour | 80.2 | 1.99 | RIKEN-GODA9AA, RIKEN-GODA9BA | To the Experiment |
| 5 | Zeatin Treated at 1 Hour | 80.67 | 2.4 | RIKEN-GODA11A, RIKEN-GODA11B | To the Experiment |
| 6 | Control at 3 Hours | 65.45 | 7.44 | RIKEN-GODA17AA, RIKEN-GODA17BA | To the Experiment |
| 6 | Zeatin Treated at 3 Hours | 67.99 | 1.6 | RIKEN-GODA19A, RIKEN-GODA19B | To the Experiment |
| 7 | Control at 30 Minutes | 72.83 | 14.07 | RIKEN-GODA1AA, RIKEN-GODA1BB | To the Experiment |
| 7 | IAA Treated at 30 Minutes | 72.88 | 10.38 | RIKEN-GODA2A, RIKEN-GODA2B | To the Experiment |
| 8 | Control at 1 Hour | 80.2 | 1.99 | RIKEN-GODA9AA, RIKEN-GODA9BA | To the Experiment |
| 8 | IAA Treated at 1 Hour | 60.77 | 9.04 | RIKEN-GODA10A, RIKEN-GODA10B | To the Experiment |
| 9 | Control at 3 Hours | 65.45 | 7.44 | RIKEN-GODA17AA, RIKEN-GODA17BA | To the Experiment |
| 9 | IAA Treated at 3 Hours | 76.75 | 15.54 | RIKEN-GODA18A, RIKEN-GODA18B | To the Experiment |
| 10 | Control at 30 Minutes | 72.83 | 14.07 | RIKEN-GODA1AA, RIKEN-GODA1BB | To the Experiment |
| 10 | ABA Treated at 30 Minutes | 66.92 | 12.73 | RIKEN-GODA5A, RIKEN-GODA5B | To the Experiment |
| 11 | Control at 1 Hour | 80.2 | 1.99 | RIKEN-GODA9AA, RIKEN-GODA9BA | To the Experiment |
| 11 | ABA Treated at 1 Hour | 70.27 | 4.58 | RIKEN-GODA13A, RIKEN-GODA13B | To the Experiment |
| 12 | Control at 3 Hours | 65.45 | 7.44 | RIKEN-GODA17AA, RIKEN-GODA17BA | To the Experiment |
| 12 | ABA Treated at 3 Hours | 72.36 | 4.97 | RIKEN-GODA21A, RIKEN-GODA21B | To the Experiment |
| 13 | Control at 30 Minutes | 72.83 | 14.07 | RIKEN-GODA1AA, RIKEN-GODA1BB | To the Experiment |
| 13 | MJ Treated at 30 Minutes | 52.75 | 11.58 | RIKEN-GODA6A, RIKEN-GODA6B | To the Experiment |
| 14 | Control at 1 Hour | 80.2 | 1.99 | RIKEN-GODA9AA, RIKEN-GODA9BA | To the Experiment |
| 14 | MJ Treated at 1 Hour | 70.03 | 4.01 | RIKEN-GODA14A, RIKEN-GODA14B | To the Experiment |
| 15 | Control at 3 Hours | 65.45 | 7.44 | RIKEN-GODA17AA, RIKEN-GODA17BA | To the Experiment |
| 15 | MJ Treated at 3 Hours | 66.17 | 8.36 | RIKEN-GODA22A, RIKEN-GODA22B | To the Experiment |
| 16 | Control at 30 Minutes | 72.83 | 14.07 | RIKEN-GODA1AA, RIKEN-GODA1BB | To the Experiment |
| 16 | GA-3 Treated at 30 Minutes | 71.78 | 6.1 | RIKEN-GODA4A, RIKEN-GODA4B | To the Experiment |
| 17 | Control at 1 Hour | 80.2 | 1.99 | RIKEN-GODA9AA, RIKEN-GODA9BA | To the Experiment |
| 17 | GA-3 Treated at 1 Hour | 74.77 | 6.97 | RIKEN-GODA12A, RIKEN-GODA12B | To the Experiment |
| 18 | Control at 3 Hours | 65.45 | 7.44 | RIKEN-GODA17AA, RIKEN-GODA17BA | To the Experiment |
| 18 | GA-3 Treated at 3 Hours | 66.28 | 10.6 | RIKEN-GODA20A, RIKEN-GODA20B | To the Experiment |
| 19 | Control at 30 Minutes | 60.53 | 2.34 | RIKEN-GODA25A, RIKEN-GODA25B | To the Experiment |
| 19 | GA-3 Treated Mutant at 30 Minutes | 72.91 | 12.28 | RIKEN-GODA26A, RIKEN-GODA26B | To the Experiment |
| 20 | Control at 1 Hour | 73.67 | 0.2 | RIKEN-GODA27A, RIKEN-GODA27B | To the Experiment |
| 20 | GA-3 Treated Mutant at 1 Hour | 77.85 | 3.35 | RIKEN-GODA28A, RIKEN-GODA28B | To the Experiment |
| 21 | Control at 3 Hours | 87.8 | 24.37 | RIKEN-GODA29A, RIKEN-GODA29B | To the Experiment |
| 21 | GA-3 Treated Mutant at 3 Hours | 69.44 | 2.93 | RIKEN-GODA30A, RIKEN-GODA30B | To the Experiment |
| 22 | Control at 30 Minutes | 72.83 | 14.07 | RIKEN-GODA1AA, RIKEN-GODA1BB | To the Experiment |
| 22 | BL Treated at 30 Minutes | 61.38 | 10.95 | RIKEN-GODA8A, RIKEN-GODA8B | To the Experiment |
| 23 | BL Control at 1 Hour | 80.2 | 1.99 | RIKEN-GODA9AA, RIKEN-GODA9BA | To the Experiment |
| 23 | Treated at 1 Hour | 67.66 | 0.79 | RIKEN-GODA16A, RIKEN-GODA16B | To the Experiment |
| 24 | Control at 3 Hours | 65.45 | 7.44 | RIKEN-GODA17AA, RIKEN-GODA17BA | To the Experiment |
| 24 | BL Treated at 3 Hours | 66.16 | 4.19 | RIKEN-GODA24A, RIKEN-GODA24B | To the Experiment |
| 25 | Control at 30 Minutes | 80.34 | 4.77 | RIKEN-GODA31A, RIKEN-GODA31B | To the Experiment |
| 25 | BL Treated Mutant at 30 Minutes | 72.28 | 2.91 | RIKEN-GODA32A, RIKEN-GODA32B | To the Experiment |
| 26 | Control at 1 Hour | 73.66 | 6.75 | RIKEN-GODA33A, RIKEN-GODA33B | To the Experiment |
| 26 | BL Treated Mutant at 1 Hour | 77.2 | 5.19 | RIKEN-GODA34A, RIKEN-GODA34B | To the Experiment |
| 27 | Control at 3 Hours | 77.55 | 3.38 | RIKEN-GODA35A, RIKEN-GODA35B | To the Experiment |
| 27 | BL Treated Mutant at 3 Hours | 64.16 | 1.12 | RIKEN-GODA36A, RIKEN-GODA36B | To the Experiment |
| 28 | Control | 64.77 | 16.07 | RIKEN-GODA1A-6, RIKEN-GODA1B-6 | To the Experiment |
| 28 | campestanol Treated | 59.76 | 11.67 | RIKEN-GODA2A-6, RIKEN-GODA2B-6 | To the Experiment |
| 28 | 6-deoxocathasterone Treated | 69.6 | 0.34 | RIKEN-GODA3A-6, RIKEN-GODA3B-6 | To the Experiment |
| 28 | cathasterone Treated | 69.16 | 4.74 | RIKEN-GODA4A-6, RIKEN-GODA4B-6 | To the Experiment |
| 28 | 6-deoxoteasterone Treated | 64.27 | 5.65 | RIKEN-GODA5A-6, RIKEN-GODA5B-6 | To the Experiment |
| 28 | teasterone Treated | 72.86 | 24.66 | RIKEN-GODA6A-6, RIKEN-GODA6B-6 | To the Experiment |
| 28 | 3-dehydro-6-deoxoteasterone Treated | 47.2 | 12.86 | RIKEN-GODA7A-6, RIKEN-GODA7B-6 | To the Experiment |
| 28 | 3-dehydroteasterone Treated | 63.49 | 6.45 | RIKEN-GODA8A-6, RIKEN-GODA8B-6 | To the Experiment |
| 28 | -deoxotyphasterol Treated | 68.92 | 1.66 | RIKEN-GODA9A-6, RIKEN-GODA9B-6 | To the Experiment |
| 28 | typhasterol Treated | 69.16 | 6.63 | RIKEN-GODA10A-6, RIKEN-GODA10B-6 | To the Experiment |
| 28 | 6-deoxocastasterone Treated | 59.57 | 0.31 | RIKEN-GODA11A-6, RIKEN-GODA11B-6 | To the Experiment |
| 28 | castasterone Treated | 63.41 | 4.86 | RIKEN-GODA12A-6, RIKEN-GODA12B-6 | To the Experiment |
| 28 | brassinolide Treated | 63.45 | 16.0 | RIKEN-GODA13A-6, RIKEN-GODA13B-6 | To the Experiment |
| 29 | Control Wild-Type | 57.88 | 2.97 | NO.10, NO.11, NO.12 | To the Experiment |
| 29 | t-zeatin Treated Wild-Type | 65.23 | 7.4 | NO.19-2, NO.20-2, NO.21-2 | To the Experiment |
| 30 | Control ARR22 Overexpressed | 60.0 | 4.59 | NO.28, NO.29, NO.30 | To the Experiment |
| 30 | t-zeatin Treated ARR22 Overexpressed | 58.13 | 3.33 | NO.31, NO.32, NO.33 | To the Experiment |
| 31 | No Treatment | 165.26 | 2.46 | RIKEN-NAKABAYASHI1A, RIKEN-NAKABAYASHI1B | To the Experiment |
| 31 | Treated with Water | 99.98 | 0.74 | RIKEN-NAKABAYASHI2A, RIKEN-NAKABAYASHI2B | To the Experiment |
| 31 | Treated with 3uM ABA | 93.74 | 1.97 | RIKEN-NAKABAYASHI3A, RIKEN-NAKABAYASHI5B | To the Experiment |
| 31 | Treated with 30uM ABA | 112.24 | 6.52 | RIKEN-NAKABAYASHI4A, RIKEN-NAKABAYASHI4B | To the Experiment |
| 32 | Control at 3 Hours | 171.22 | 22.97 | RIKEN-LI1A, RIKEN-LI1B | To the Experiment |
| 32 | Giberellin Treated at 3 Hours | 135.53 | 9.38 | RIKEN-LI4A, RIKEN-LI4B | To the Experiment |
| 33 | Control at 6 Hours | 152.59 | 2.2 | RIKEN-LI2A, RIKEN-LI2B | To the Experiment |
| 33 | Giberellin Treated at 6 Hours | 155.47 | 2.52 | RIKEN-LI5A, RIKEN-LI5B | To the Experiment |
| 34 | Control at 9 Hours | 134.41 | 22.15 | RIKEN-LI3A, RIKEN-LI3B | To the Experiment |
| 34 | Giberellin Treated at 9 Hours | 135.25 | 66.93 | RIKEN-LI6A, RIKEN-LI6B | To the Experiment |

**Suppl. Table S1.**
Gene expression data for the *MDF* gene regulation in response to hormones obtained from https://bar.utoronto.ca/efp/cgi-bin/efpWeb.cgi?dataSource=Hormone&mode=Absolute&primaryGene=At5g16780&secondaryGene=At3g27340&override=&threshold=604.48&modeMask_low=None&modeMask_stddev=None).
