## Supplementary material for "MDF regulates a network of auxin-dependent and -independent pathways of adventitious root regeneration in *Arabidopsis*": Table S2

| Group # | Tissue | Expression Level | Standard Deviation | Samples | Links |
|---|---|---|---|---|---|
| 1 | Control Shoot 0 Hour | 82.94 | 10.78 | AtGen_6_0011, AtGen_6_0012 | To the Experiment |
| 1 | Cold Shoot 0 Hour | 82.94 | 10.78 | AtGen_6_0011, AtGen_6_0012 | To the Experiment |
| 1 | Osmotic Shoot 0 Hour | 82.94 | 10.78 | AtGen_6_0011, AtGen_6_0012 | To the Experiment |
| 1 | Salt Shoot 0 Hour | 82.94 | 10.78 | AtGen_6_0011, AtGen_6_0012 | To the Experiment |
| 1 | Drought Shoot 0 Hour | 82.94 | 10.78 | AtGen_6_0011, AtGen_6_0012 | To the Experiment |
| 1 | Genotoxic Shoot 0 Hour | 82.94 | 10.78 | AtGen_6_0011, AtGen_6_0012 | To the Experiment |
| 1 | Oxidative Shoot 0 Hour | 82.94 | 10.78 | AtGen_6_0011, AtGen_6_0012 | To the Experiment |
| 1 | UV-B Shoot 0 Hour | 82.94 | 10.78 | AtGen_6_0011, AtGen_6_0012 | To the Experiment |
| 1 | Wounding Shoot 0 Hour | 82.94 | 10.78 | AtGen_6_0011, AtGen_6_0012 | To the Experiment |
| 1 | Heat Shoot 0 Hour | 82.94 | 10.78 | AtGen_6_0011, AtGen_6_0012 | To the Experiment |
| 2 | Control Root 0 Hour | 105.02 | 0.59 | AtGen_6_0021, AtGen_6_0022 | To the Experiment |
| 2 | Cold Root 0 Hour | 105.02 | 0.59 | AtGen_6_0021, AtGen_6_0022 | To the Experiment |
| 2 | Osmotic Root 0 Hour | 105.02 | 0.59 | AtGen_6_0021, AtGen_6_0022 | To the Experiment |
| 2 | Salt Root 0 Hour | 105.02 | 0.59 | AtGen_6_0021, AtGen_6_0022 | To the Experiment |
| 2 | Drought Root 0 Hour | 105.02 | 0.59 | AtGen_6_0021, AtGen_6_0022 | To the Experiment |
| 2 | Genotoxic Root 0 Hour | 105.02 | 0.59 | AtGen_6_0021, AtGen_6_0022 | To the Experiment |
| 2 | Oxidative Root 0 Hour | 105.02 | 0.59 | AtGen_6_0021, AtGen_6_0022 | To the Experiment |
| 2 | UV-B Root 0 Hour | 105.02 | 0.59 | AtGen_6_0021, AtGen_6_0022 | To the Experiment |
| 2 | Wounding Root 0 Hour | 105.02 | 0.59 | AtGen_6_0021, AtGen_6_0022 | To the Experiment |
| 2 | Heat Root 0 Hour | 105.02 | 0.59 | AtGen_6_0021, AtGen_6_0022 | To the Experiment |
| 3 | Control Shoot After 15 Minutes | 76.45 | 1.75 | AtGen_6_0711, AtGen_6_0712 | To the Experiment |
| 3 | Drought Shoot After 15 Minutes | 84.7 | 7.64 | AtGen_6_4711, AtGen_6_4712 | To the Experiment |
| 3 | UV-B Shoot After 15 Minutes | 84.5 | 17.44 | AtGen_6_7711, AtGen_6_7712 | To the Experiment |
| 3 | Wounding Shoot After 15 Minutes | 85.19 | 5.28 | AtGen_6_8715, AtGen_6_8712 | To the Experiment |
| 3 | Heat Shoot After 15 Minutes | 86.44 | 11.79 | AtGen_6_9711, AtGen_6_9712 | To the Experiment |
| 4 | Control Root After 15 Minutes | 90.44 | 4.83 | AtGen_6_0721, AtGen_6_0722 | To the Experiment |
| 4 | Drought Root After 15 Minutes | 96.25 | 6.77 | AtGen_6_4721, AtGen_6_4722 | To the Experiment |
| 4 | UV-B Root After 15 Minutes | 105.0 | 1.89 | AtGen_6_7721, AtGen_6_7722 | To the Experiment |
| 4 | Wounding Root After 15 Minutes | 104.38 | 3.25 | AtGen_6_8723, AtGen_6_8724 | To the Experiment |
| 4 | Heat Root After 15 Minutes | 94.22 | 0.96 | AtGen_6_9721, AtGen_6_9722 | To the Experiment |
| 5 | Control Shoot After 30 Minutes | 98.4 | 12.64 | AtGen_6_0111, AtGen_6_0112 | To the Experiment |
| 5 | Cold Shoot After 30 Minutes | 70.03 | 1.26 | AtGen_6_1111, AtGen_6_1112 | To the Experiment |
| 5 | Osmotic Shoot After 30 Minutes | 70.67 | 5.12 | AtGen_6_2111, AtGen_6_2112 | To the Experiment |
| 5 | Salt Shoot After 30 Minutes | 85.02 | 7.51 | AtGen_6_3111, AtGen_6_3112 | To the Experiment |
| 5 | Drought Shoot After 30 Minutes | 78.33 | 6.38 | AtGen_6_4111, AtGen_6_4112 | To the Experiment |
| 5 | Genotoxic Shoot After 30 Minutes | 110.11 | 7.84 | AtGen_6_5111, AtGen_6_5112 | To the Experiment |
| 5 | Oxidative Shoot After 30 Minutes | 98.47 | 3.06 | AtGen_6_6111, AtGen_6_6112 | To the Experiment |
| 5 | UV-B Shoot After 30 Minutes | 82.28 | 1.0 | AtGen_6_7111, AtGen_6_7112 | To the Experiment |
| 5 | Wounding Shoot After 30 Minutes | 80.5 | 0.49 | AtGen_6_8111, AtGen_6_8112 | To the Experiment |
| 5 | Heat Shoot After 30 Minutes | 88.79 | 3.03 | AtGen_6_9111, AtGen_6_9112 | To the Experiment |
| 6 | Control Root After 30 Minutes | 108.19 | 7.35 | AtGen_6_0121, AtGen_6_0122 | To the Experiment |
| 6 | Cold Root After 30 Minutes | 82.72 | 9.72 | AtGen_6_1121, AtGen_6_1122 | To the Experiment |
| 6 | Osmotic Root After 30 Minutes | 88.81 | 5.69 | AtGen_6_2121, AtGen_6_2122 | To the Experiment |
| 6 | Salt Root After 30 Minutes | 112.46 | 4.2 | AtGen_6_3121, AtGen_6_3122 | To the Experiment |
| 6 | Drought Root After 30 Minutes | 93.44 | 2.13 | AtGen_6_4121, AtGen_6_4122 | To the Experiment |
| 6 | Genotoxic Root After 30 Minutes | 103.85 | 0.04 | AtGen_6_5121, AtGen_6_5122 | To the Experiment |
| 6 | Oxidative Root After 30 Minutes | 102.25 | 4.99 | AtGen_6_6122, AtGen_6_6124 | To the Experiment |
| 6 | UV-B Root After 30 Minutes | 108.04 | 2.55 | AtGen_6_7121, AtGen_6_7122 | To the Experiment |
| 6 | Wounding Root After 30 Minutes | 97.33 | 0.59 | AtGen_6_8124, AtGen_6_8126 | To the Experiment |
| 6 | Heat Root After 30 Minutes | 119.44 | 17.3 | AtGen_6_9121, AtGen_6_9122 | To the Experiment |
| 7 | Control Shoot After 1 Hour | 87.42 | 11.74 | AtGen_6_0211, AtGen_6_0212 | To the Experiment |
| 7 | Cold Shoot After 1 Hour | 69.44 | 5.71 | AtGen_6_1211, AtGen_6_1212 | To the Experiment |
| 7 | Osmotic Shoot After 1 Hour | 80.83 | 0.02 | AtGen_6_2211, AtGen_6_2212 | To the Experiment |
| 7 | Salt Shoot After 1 Hour | 109.25 | 4.86 | AtGen_6_3211, AtGen_6_3212 | To the Experiment |
| 7 | Drought Shoot After 1 Hour | 80.37 | 1.09 | AtGen_6_4211, AtGen_6_4212 | To the Experiment |
| 7 | Genotoxic Shoot After 1 Hour | 110.86 | 4.11 | AtGen_6_5211, AtGen_6_5212 | To the Experiment |
| 7 | Oxidative Shoot After 1 Hour | 84.97 | 6.94 | AtGen_6_6211, AtGen_6_6212 | To the Experiment |
| 7 | UV-B Shoot After 1 Hour | 71.42 | 17.12 | AtGen_6_7211, AtGen_6_7212 | To the Experiment |
| 7 | Wounding Shoot After 1 Hour | 82.03 | 6.97 | AtGen_6_8211, AtGen_6_8214 | To the Experiment |
| 7 | Heat Shoot After 1 Hour | 80.06 | 8.61 | AtGen_6_9211, AtGen_6_9212 | To the Experiment |
| 8 | Control Root After 1 Hour | 93.31 | 3.13 | AtGen_6_0221, AtGen_6_0222 | To the Experiment |
| 8 | Cold Root After 1 Hour | 77.69 | 3.55 | AtGen_6_1221, AtGen_6_1222 | To the Experiment |
| 8 | Osmotic Root After 1 Hour | 98.59 | 21.02 | AtGen_6_2221, AtGen_6_2222 | To the Experiment |
| 8 | Salt Root After 1 Hour | 105.11 | 0.64 | AtGen_6_3221, AtGen_6_3222 | To the Experiment |
| 8 | Drought Root After 1 Hour | 97.41 | 8.28 | AtGen_6_4221, AtGen_6_4222 | To the Experiment |
| 8 | Genotoxic Root After 1 Hour | 107.58 | 24.37 | AtGen_6_5221, AtGen_6_5222 | To the Experiment |
| 8 | Oxidative Root After 1 Hour | 108.47 | 6.96 | AtGen_6_6223, AtGen_6_6224 | To the Experiment |
| 8 | UV-B Root After 1 Hour | 112.82 | 1.53 | AtGen_6_7221, AtGen_6_7222 | To the Experiment |
| 8 | Wounding Root After 1 Hour | 105.07 | 3.03 | AtGen_6_8224, AtGen_6_8225 | To the Experiment |
| 8 | Heat Root After 1 Hour | 100.64 | 3.97 | AtGen_6_9221, AtGen_6_9222 | To the Experiment |
| 9 | Control Shoot After 3 Hours | 95.56 | 7.59 | AtGen_6_0311, AtGen_6_0312 | To the Experiment |
| 9 | Cold Shoot After 3 Hours | 76.24 | 2.09 | AtGen_6_1311, AtGen_6_1312 | To the Experiment |
| 9 | Osmotic Shoot After 3 Hours | 80.61 | 3.04 | AtGen_6_2311, AtGen_6_2312 | To the Experiment |
| 9 | Salt Shoot After 3 Hours | 90.83 | 15.08 | AtGen_6_3311, AtGen_6_3312 | To the Experiment |
| 9 | Drought Shoot After 3 Hours | 92.41 | 4.21 | AtGen_6_4311, AtGen_6_4312 | To the Experiment |
| 9 | Genotoxic Shoot After 3 Hours | 106.65 | 7.23 | AtGen_6_5311, AtGen_6_5312 | To the Experiment |
| 9 | Oxidative Shoot After 3 Hours | 103.34 | 5.53 | AtGen_6_6311, AtGen_6_6312 | To the Experiment |
| 9 | UV-B Shoot After 3 Hours | 103.54 | 2.62 | AtGen_6_7311, AtGen_6_7312 | To the Experiment |
| 9 | Wounding Shoot After 3 Hours | 87.29 | 4.74 | AtGen_6_8313, AtGen_6_8314 | To the Experiment |
| 9 | Heat Shoot After 3 Hours | 97.78 | 7.66 | AtGen_6_9311, AtGen_6_9312 | To the Experiment |
| 10 | Control Root After 3 Hours | 108.89 | 7.59 | AtGen_6_0321, AtGen_6_0322 | To the Experiment |
| 10 | Cold Root After 3 Hours | 84.86 | 10.47 | AtGen_6_1321, AtGen_6_1322 | To the Experiment |
| 10 | Osmotic Root After 3 Hours | 123.81 | 2.62 | AtGen_6_2321, AtGen_6_2322 | To the Experiment |
| 10 | Salt Root After 3 Hours | 86.47 | 8.73 | AtGen_6_3321, AtGen_6_3322 | To the Experiment |
| 10 | Drought Root After 3 Hours | 99.97 | 2.41 | AtGen_6_4321, AtGen_6_4322 | To the Experiment |
| 10 | Genotoxic Root After 3 Hours | 123.34 | 2.31 | AtGen_6_5321, AtGen_6_5322 | To the Experiment |
| 10 | Oxidative Root After 3 Hours | 113.88 | 23.83 | AtGen_6_6322, AtGen_6_6323 | To the Experiment |
| 10 | UV-B Root After 3 Hours | 113.72 | 5.2 | AtGen_6_7321, AtGen_6_7322 | To the Experiment |
| 10 | Wounding Root After 3 Hours | 117.94 | 5.48 | AtGen_6_8324, AtGen_6_8325 | To the Experiment |
| 10 | Heat Root After 3 Hours | 124.59 | 2.09 | AtGen_6_9321, AtGen_6_9322 | To the Experiment |
| 11 | Control Shoot After 4 Hours | 81.81 | 0.57 | AtGen_6_0811, AtGen_6_0812 | To the Experiment |
| 11 | Heat Shoot After 4 Hours | 120.56 | 2.18 | AtGen_6_9811, AtGen_6_9812 | To the Experiment |
| 12 | Control Root After 4 Hours | 116.53 | 6.48 | AtGen_6_0821, AtGen_6_0822 | To the Experiment |
| 12 | Heat Root After 4 Hours | 153.56 | 9.89 | AtGen_6_9821, AtGen_6_9822 | To the Experiment |
| 13 | Control Shoot After 6 Hours | 77.1 | 9.32 | AtGen_6_0411, AtGen_6_0412 | To the Experiment |
| 13 | Cold Shoot After 6 Hours | 82.23 | 21.28 | AtGen_6_1411, AtGen_6_1412 | To the Experiment |
| 13 | Osmotic Shoot After 6 Hours | 107.05 | 11.55 | AtGen_6_2411, AtGen_6_2412 | To the Experiment |
| 13 | Salt Shoot After 6 Hours | 79.92 | 1.43 | AtGen_6_3411, AtGen_6_3412 | To the Experiment |
| 13 | Drought Shoot After 6 Hours | 108.49 | 2.02 | AtGen_6_4411, AtGen_6_4412 | To the Experiment |
| 13 | Genotoxic Shoot After 6 Hours | 105.37 | 11.06 | AtGen_6_5411, AtGen_6_5412 | To the Experiment |
| 13 | Oxidative Shoot After 6 Hours | 104.81 | 5.12 | AtGen_6_6411, AtGen_6_6412 | To the Experiment |
| 13 | UV-B Shoot After 6 Hours | 110.31 | 14.57 | AtGen_6_7411, AtGen_6_7412 | To the Experiment |
| 13 | Wounding Shoot After 6 Hours | 94.22 | 14.8 | AtGen_6_8411, AtGen_6_8412 | To the Experiment |
| 13 | Heat Shoot After 6 Hours | 92.06 | 4.52 | AtGen_6_9411, AtGen_6_9412 | To the Experiment |
| 14 | Control Root After 6 Hours | 113.06 | 3.13 | AtGen_6_0421, AtGen_6_0422 | To the Experiment |
| 14 | Cold Root After 6 Hours | 100.78 | 1.95 | AtGen_6_1421, AtGen_6_1422 | To the Experiment |
| 14 | Osmotic Root After 6 Hours | 124.05 | 7.1 | AtGen_6_2421, AtGen_6_2422 | To the Experiment |
| 14 | Salt Root After 6 Hours | 67.5 | 6.45 | AtGen_6_3421, AtGen_6_3422 | To the Experiment |
| 14 | Drought Root After 6 Hours | 131.58 | 10.19 | AtGen_6_4421, AtGen_6_4422 | To the Experiment |
| 14 | Genotoxic Root After 6 Hours | 127.91 | 11.39 | AtGen_6_5421, AtGen_6_5422 | To the Experiment |
| 14 | Oxidative Root After 6 Hours | 113.33 | 0.82 | AtGen_6_6421, AtGen_6_6422 | To the Experiment |
| 14 | UV-B Root After 6 Hours | 121.97 | 0.98 | AtGen_6_7421, AtGen_6_7422 | To the Experiment |
| 14 | Wounding Root After 6 Hours | 123.8 | 18.6 | AtGen_6_8423, AtGen_6_8424 | To the Experiment |
| 14 | Heat Root After 6 Hours | 134.25 | 7.76 | AtGen_6_9421, AtGen_6_9422 | To the Experiment |
| 15 | Control Shoot After 12 Hours | 79.09 | 19.17 | AtGen_6_0511, AtGen_6_0512 | To the Experiment |
| 15 | Cold Shoot After 12 Hours | 112.71 | 8.92 | AtGen_6_1511, AtGen_6_1512 | To the Experiment |
| 15 | Osmotic Shoot After 12 Hours | 88.84 | 7.65 | AtGen_6_2511, AtGen_6_2512 | To the Experiment |
| 15 | Salt Shoot After 12 Hours | 85.75 | 1.83 | AtGen_6_3511, AtGen_6_3512 | To the Experiment |
| 15 | Drought Shoot After 12 Hours | 83.72 | 11.39 | AtGen_6_4511, AtGen_6_4512 | To the Experiment |
| 15 | Genotoxic Shoot After 12 Hours | 99.12 | 0.82 | AtGen_6_5511, AtGen_6_5512 | To the Experiment |
| 15 | Oxidative Shoot After 12 Hours | 93.45 | 10.09 | AtGen_6_6511, AtGen_6_6512 | To the Experiment |
| 15 | UV-B Shoot After 12 Hours | 106.93 | 8.56 | AtGen_6_7511, AtGen_6_7512 | To the Experiment |
| 15 | Wounding Shoot After 12 Hours | 84.32 | 7.2 | AtGen_6_8511, AtGen_6_8512 | To the Experiment |
| 15 | Heat Shoot After 12 Hours | 76.52 | 4.21 | AtGen_6_9511, AtGen_6_9512 | To the Experiment |
| 16 | Control Root After 12 Hours | 118.05 | 5.37 | AtGen_6_0521, AtGen_6_0522 | To the Experiment |
| 16 | Cold Root After 12 Hours | 121.0 | 1.7 | AtGen_6_1521, AtGen_6_1522 | To the Experiment |
| 16 | Osmotic Root After 12 Hours | 131.75 | 18.58 | AtGen_6_2521, AtGen_6_2522 | To the Experiment |
| 16 | Salt Root After 12 Hours | 87.81 | 32.24 | AtGen_6_3521, AtGen_6_3522 | To the Experiment |
| 16 | Drought Root After 12 Hours | 122.36 | 20.73 | AtGen_6_4521, AtGen_6_4522 | To the Experiment |
| 16 | Genotoxic Root After 12 Hours | 110.19 | 9.36 | AtGen_6_5521, AtGen_6_5522 | To the Experiment |
| 16 | Oxidative Root After 12 Hours | 109.23 | 0.81 | AtGen_6_6523, AtGen_6_6524 | To the Experiment |
| 16 | UV-B Root After 12 Hours | 118.03 | 5.64 | AtGen_6_7521, AtGen_6_7522 | To the Experiment |
| 16 | Wounding Root After 12 Hours | 109.74 | 16.28 | AtGen_6_8524, AtGen_6_8525 | To the Experiment |
| 16 | Heat Root After 12 Hours | 126.76 | 0.89 | AtGen_6_9521, AtGen_6_9522 | To the Experiment |
| 17 | Control Shoot After 24 Hours | 99.17 | 7.92 | AtGen_6_0611, AtGen_6_0612 | To the Experiment |
| 17 | Cold Shoot After 24 Hours | 123.47 | 12.3 | AtGen_6_1611, AtGen_6_1612 | To the Experiment |
| 17 | Osmotic Shoot After 24 Hours | 83.69 | 5.15 | AtGen_6_2611, AtGen_6_2612 | To the Experiment |
| 17 | Salt Shoot After 24 Hours | 79.25 | 4.95 | AtGen_6_3611, AtGen_6_3612 | To the Experiment |
| 17 | Drought Shoot After 24 Hours | 91.92 | 7.47 | AtGen_6_4611, AtGen_6_4612 | To the Experiment |
| 17 | Genotoxic Shoot After 24 Hours | 89.67 | 0.66 | AtGen_6_5611, AtGen_6_5612 | To the Experiment |
| 17 | Oxidative Shoot After 24 Hours | 94.91 | 0.22 | AtGen_6_6611, AtGen_6_6612 | To the Experiment |
| 17 | UV-B Shoot After 24 Hours | 95.23 | 15.49 | AtGen_6_7611, AtGen_6_7612 | To the Experiment |
| 17 | Wounding Shoot After 24 Hours | 94.42 | 0.27 | AtGen_6_8611, AtGen_6_8612 | To the Experiment |
| 17 | Heat Shoot After 24 Hours | 88.05 | 4.79 | AtGen_6_9611, AtGen_6_9612 | To the Experiment |
| 18 | Control Root After 24 Hours | 107.55 | 10.66 | AtGen_6_0621, AtGen_6_0622 | To the Experiment |
| 18 | Cold Root After 24 Hours | 120.73 | 1.39 | AtGen_6_1621, AtGen_6_1622 | To the Experiment |
| 18 | Osmotic Root After 24 Hours | 95.01 | 15.97 | AtGen_6_2621, AtGen_6_2622 | To the Experiment |
| 18 | Salt Root After 24 Hours | 84.74 | 10.38 | AtGen_6_3621, AtGen_6_3622 | To the Experiment |
| 18 | Drought Root After 24 Hours | 112.63 | 10.87 | AtGen_6_4621, AtGen_6_4622 | To the Experiment |
| 18 | Genotoxic Root After 24 Hours | 106.91 | 10.75 | AtGen_6_5621, AtGen_6_5622 | To the Experiment |
| 18 | Oxidative Root After 24 Hours | 129.24 | 34.0 | AtGen_6_6621, AtGen_6_6625 | To the Experiment |
| 18 | UV-B Root After 24 Hours | 110.16 | 12.4 | AtGen_6_7621, AtGen_6_7622 | To the Experiment |
| 18 | Wounding Root After 24 Hours | 105.84 | 10.21 | AtGen_6_8621, AtGen_6_8622 | To the Experiment |
| 18 | Heat Root After 24 Hours | 106.41 | 4.84 | AtGen_6_9621, AtGen_6_9622 | To the Experiment |

**Suppl. Table S2.**

Gene expression data for the *MDF* gene regulation in response to abiotic stresses obtained from https://bar.utoronto.ca/ efp/cgi-bin/efpWeb.cgi?dataSource=Hormone&mode=Absolute&primaryGene=At5g16780&secondaryGene=At3g27340&override=&threshold=604.48&modeMask_low=None&modeMask_stddev=None).
