## Supplementary material for "MDF regulates a network of auxin-dependent and -independent pathways of adventitious root regeneration in *Arabidopsis*": Table S3

**Table S3. PCR Primers**

| Primer name | Sequence 5'-3' |
|---|---|
| **Cloning:** | |
| pDONR207 Forward | TCGCGTTAACGCTAGCATGGATCTC |
| pDONR207 Reverse | GTAACATCAGAGATTTTGAGACAC |
| *RAP2.7* Forward | GGGGACAAGTTTGTACAAAAAAGCAGGCTTAATGTTGGATCTTAACCTCAACGCT |
| *RAP2.7* Reverse | GGGGACCACTTTGTACAAGAAAGCTGGGTTTAAGGGTGTGGATAAAAGTAACCACGT |
| *NAC1* Forward | GGGGACAAGTTTGTACAAAAAAGCAGGCTTAATGGAGACGGAAGAAGAGATGAAG |
| *NAC1* Reverse | GGGGACCACTTTGTACAAGAAAGCTGGGTTTCAGCAATTCCAAACAGTGCTTGGA |
| **RT-qPCR:** | |
| *UBC* Forward | CTGCGACTCAGGGATCTTCTAA |
| *UBC* Reverse | TTGTGCCATTGAATTGAACCC |
| *UBQ10* Forward | GGCCTTGTATAATCCCTGATGAATAAG |
| *UBQ10* Reverse | AAAGAGATAACAGGAACGGAAACATAGT |
| *ACT2* Forward | CTTGCACCAAGCAGCATGAA |
| *ACT2* Reverse | CCGATCCAGACACTGTACTTCCTT |
| *MDF* Forward | GGCCTGGAAAAATGAAGGA |
| *MDF* Reverse | GGCCACTGAGGACAAGGTAA |
| *PIN1* Forward | TCAGGGGAATAGTAACGACAACCAG |
| *PIN1* Reverse | ATCACACTTGTTGGTGGCATCACCT |
| *PIN3* Forward | ATTCCTCTCTACGTGGCCATGATCC |
| *PIN3* Reverse | AGAGAGGAGAGGGACGGCGAAG |
| *RAP2.7* Forward | TAATGGTAGAGAAGCAGTCACGAA |
| *RAP2.7* Reverse | TGGATAAAAGTAACCACGTGTTGC |
| *YUC1* Forward | TTCATGTGTTGCCAAGGGAGATAC |

| | |
|---|---|
| *YUC1* Reverse | ACCAATTTCGCCAGCGATCTTAAC |
| *NAC1* Forward | TGGGATGAGGAAGACATTGGTTTT |
| *NAC1* Reverse | TCAATCTTAGTGAGCTGACTGAGT |
| *WOX5* Forward | AAGCTTACGTGGCAACAATAACGG |
| *WOX5* Reverse | AAGATCTAATGGCGGTGGATGTTC |
| *IAA1* forward | CGGTTAGATCTCACTGGAGGCCAT |
| *IAA1* reverse | ACTTGCTCCTCCTCCTGCAAAAAC |
| *IAA2* forward | AGGAAGAGTCTAGAGCAGGAGC |
| *IAA2* reverse | ACTGGATGTTGGTTGGTGATG |
